# Higher-order interaction of brain microstructural and functional connectome

**DOI:** 10.1101/2021.11.11.467196

**Authors:** Hao Wang, Hui-Jun Wu, Yang-Yu Liu, Linyuan Lü

**Affiliations:** Institute of Fundamental and Frontier Sciences, University of Electronic Science and Technology of China, Chengdu 611731, P. R. China.; Yangtze Delta Region Institute (Huzhou), University of Electronic Science and Technology of China, Huzhou 313001, P. R. China.; G. Oppenheimer Center for Neurobiology of Stress & Resilience, UCLA Vatche and Tamar Manoukian Division of Digestive Diseases, David Geffen School of Medicine, University of California at Los Angeles, Los Angeles, CA 90095, USA.; School of Media & Communication, Shanghai Jiao Tong University, Shanghai 200240, P. R. China.; Channing Division of Network Medicine, Brigham and Women’s Hospital and Harvard Medical School, Boston, MA 02115, USA.; Beijing Computational Science Research Center, Beijing 100193, P. R. China.

**Keywords:** brain network, complex systems, higher-order, hub, myelin

## Abstract

Despite a relatively fixed anatomical structure, the human brain can support rich cognitive functions, triggering particular interest in investigating structure-function relationships. Myelin is a vital brain microstructure marker, yet the individual microstructure-function relationship is poorly understood. Here, we explore the brain microstructure-function relationships using a higher-order framework. Global (network-level) higher-order microstructure-function relationships negatively correlate with male participants’ personality scores and decline with aging. Nodal (node-level) higher-order microstructure-function relationships are not aligned uniformly throughout the brain, being stronger in association cortices and lower in sensory cortices, showing gender differences. Notably, higher-order microstructure-function relationships are maintained from the whole-brain to local circuits, which uncovers a compelling and straightforward principle of brain structure-function interactions. Additionally, targeted artificial attacks can disrupt these higher-order relationships, and the main results are robust against several factors. Together, our results increase the collective knowledge of higher-order structure-function interactions that may underlie cognition, individual differences, and aging.

## INTRODUCTION

The human brain, unique among body organs, is a complex system (Bullmore & Sporns, 2009; Raichle et al., 2001). In this system, ever-changing human cognitive processes rely on a relatively unchanged anatomical structure, comprising about 86 billions of neurons interconnected through trillions of synapses (Landhuis, 2017). For relatively unchanged brain structural properties, myelin plays an important role that underlie behavior and learning (Bonetto et al., 2021). The formation of a myelin sheath is termed myelination, which is an essential indicator of brain maturation (Nave, 2010) and vital for neural circuit formation (Hill et al., 2018). Myelin probably shapes the functional activity (Fornari et al., 2007), correlates with psychiatric traits (Ziegler et al., 2019), and interacts with the functional connectome (Lariviere et al., 2020). Technically, myelin owns a putative microstructural MRI marker characterized by the ratio of T1- and T2-weighted (T1w/T2w) MRI images (Glasser & Van Essen, 2011; Sydnor et al., 2021), and a macroscale myelin-based microstructural brain network can be constructed based on the similarity of myelin contents between cortical regions (Wu et al., 2018), enabling the utility of myelin evidence to explore complex structure-function relationships in the human brain.

To capture the ever-changing properties of the human brain, the brain network from the resting state is an excellent entry point. Neuroimaging experiments have shown that the brain maintains a high level of activity even when it is nominally “at rest” (Raichle et al., 2001). The similarity of brain activity among multiple brain regions (Kelly et al., 2012), described as functional connectivity (FC), holds the key to understanding neurological disorders and even consciousness itself (Raichle, 2015). Therefore, clarifying the relationships between the microstructural network and the resting-state functional network is a necessary step forward to uncover the mystery of how a relatively static structure produces abundant functions (Park & Friston, 2013). Over the past decade, analyzing the structure-function relationship, under the low-order frameworks that focus on node or edge level, has dramatically expanded our comprehension of the brain functional-structural interactions, including direct edge-to-edge comparisons (Gu et al., 2021; Honey et al., 2009), multivariate statistical models (Misic et al., 2016), network-theoretic models (Vazquez-Rodriguez et al., 2019). And structure-function shows a medium level, but significant correlations (Honey et al., 2009; Mollink et al., 2019) and the nodal structure-function correspondence is not uniform across the brain (Vazquez-Rodriguez et al., 2019). Similar lower-order (edge-level) methods have also been conducted in myelin studies, constructing a structural covariance network at the population level and exploring its relationship with electrophysiological networks (Hunt et al., 2016). However, most myelin studies constructed a structural covariance network at the population level (Ma & Zhang, 2017; Melie-Garcia et al., 2018), making individual cognitive or behavior predictions impossible. Therefore, examining the myelin microstructural and functional relationship at the individual level is urgently needed but is still elusive.

Recently, higher-order representations (beyond the node or edge level) emerged, including simplicial complexes (Giusti et al., 2015), persistent homology (Liang & Wang, 2017), neural network (Suárez et al., 2021), hypergraphs (Battiston et al., 2020), subgraphs (Przulj, 2007), and motifs (Angulo et al., 2015; Benson et al., 2016), which have proven to be extremely useful in understanding and comparing complex networks. These findings reveal some higher-order connectivity patterns in brain networks (Petri et al., 2014; Sizemore et al., 2018). Nevertheless, there is still no study about the individual myelin microstructure-function relationship using higher-order representations.

Here we quantify the individual-level microstructure-function relationship using a higher-order framework. In current study, *higher-order* is defined as the interactions among orbits feature derived from subgraphs (Przulj, 2007), which is beyond the traditional pairwise interactions (Battiston et al., 2020). We use 11 non-redundant orbits because they are most efficient in computing and perform well in clustering networks with the same topological properties (Yaveroglu et al., 2014). As a comparison, a lower-order analysis is also performed.

We first reconstruct the microstructural covariance network at the group level and explore its relationship with the mean functional network from resting-state functional MRI (fMRI), using a simple multilinear model in a cohort of 198 healthy participants, to verify whether the myelin-based network can predict the functional network with a similar pattern with diffusion MRI (Vazquez-Rodriguez et al., 2019). Then, we apply a probability distribution function (PDF)-based method (Li et al., 2021) to construct the individual microstructural network and investigate its relations with the static and dynamic functional network using both lower-order and higher-order frameworks, verifying whether the higher-order relationships are different from the lower-order ones and whether the nodal higher-order relationships are uniform across the brain.

Because the individual microstructure network enables us to explore the brain-cognition relations, we link this higher-order microstructure-function relationship with individual cognitive scores and development across the three age groups. Our results reveal that the human brain owns stronger higher-order microstructure-function interactions from the whole brain to local circuits, and nodal higher-order microstructure-function relationship is not uniform in the brain, as an idiosyncratic feature of the brain, relating to cognition and demonstrating gender differences. These findings provide insight into understanding the human brain’s organization principles, its development, individual differences, and inspiring brain-like computations.

## RESULTS

We leveraged the HCP S1200 New Subjects dataset (of 213 participants, unprocessed data) (Van Essen et al., 2012) to construct the myelin-based microstructural network and resting-state functional network. After quality control, we discarded *eight* participants due to a largely framewise displacement and dropped *seven* participants with age greater than 36 due to the small sample size. Finally, we obtained quality-controlled resting-state fMRI (rfMRI) images, T1-weighted images, and T2-weighted images of 198 participants (108 males and 90 females, see ***SI Appendix,* Table S1** for a complete list of participant IDs). We constructed the structural and functional networks as follows:

### Structural network

Two types of structural networks are constructed. One is the group-level structural network, named the *microstructural covariance network*. Using Gordon parcellation (Gordon et al., 2016) with 333 regions, we extract the mean myelin content (T1w/T2w ratio) for each participant’s region, yielding a 198 × 333 matrix. For each paired region, we perform the Spearman correlation across the 198 participants, resulting in a 333 × 333 matrix, known as the covariance network (He et al., 2007), which measures the covariance of information between different regions of the brain in the population. A group-level microstructural covariance network is obtained (**Fig. 1a**). The other one is the individual-level structural network, named the *individual myelin-based microstructural networks.* For each participant, the same brain parcellation is used, then we estimate the probability distribution function (PDF) of each brain region’s myelin content. Subsequently, we calculate the similarity between the PDFs of any paired brain regions, resulting in a 333 × 333 matrix, i.e., the *individual microstructural network* (***SI Appendix***, ***Materials and Methods***).

**Figure 1.**
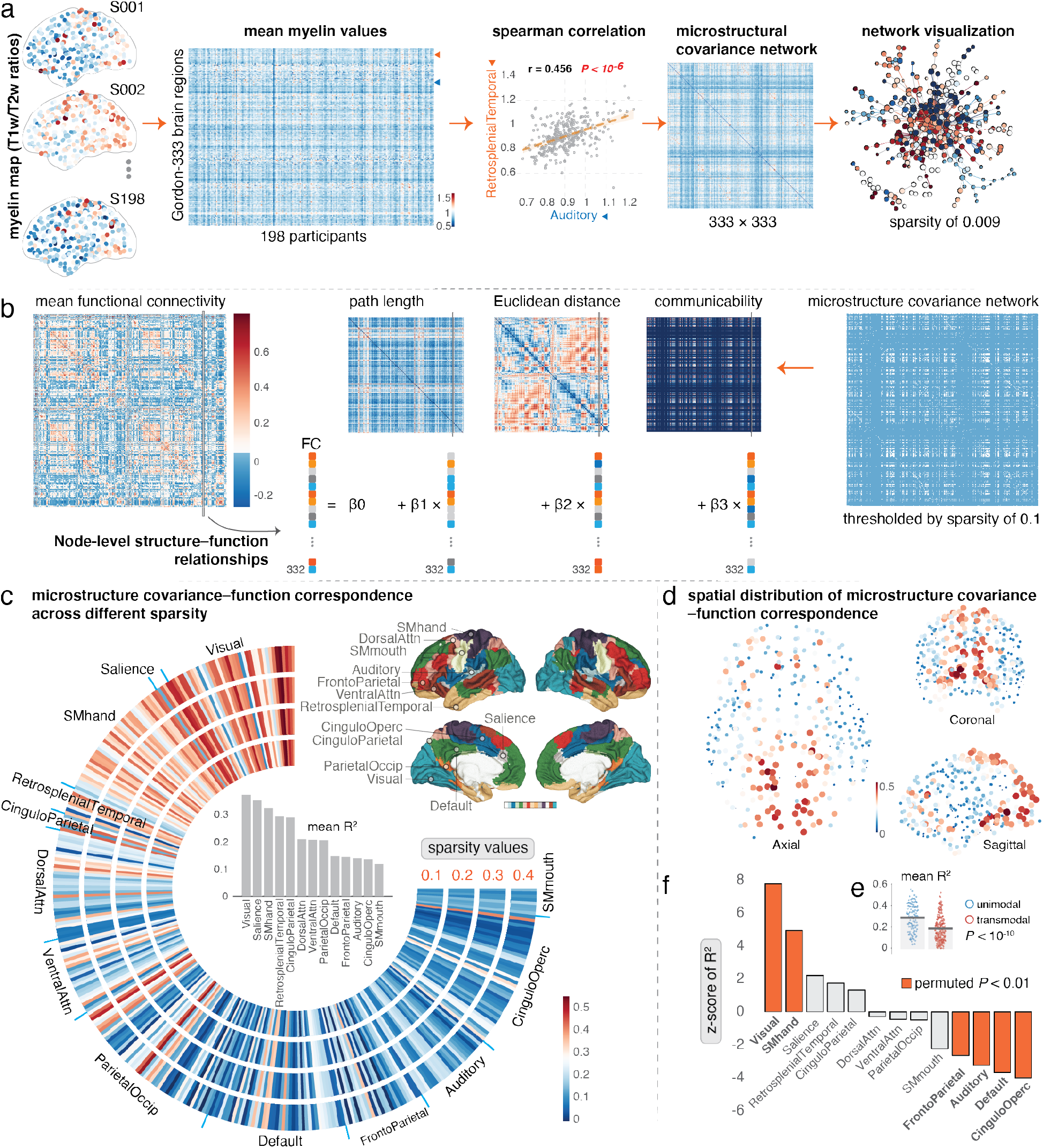
Correspondence of microstructural covariance and mean functional network. (**a**) Flowchart for constructing microstructure covariance network, using the Gordon-333 atlas to extract the mean myelin content for each region and estimate the spearman correlation for any paired regions across the population, resulting in a 333 × 333 matrix, named *microstructural covariance network*. (**b**) Node-level microstructural covariance and functional correspondence by a multilinear regression model. (**c**) Correspondence of microstructural covariance and mean functional network across different sparsity values. A similar pattern is observed for different sparsity values; the mean correspondence value is calculated for each module (13 modules in Gordon atlas). (**d**) Spatial distribution of structure-function correspondence with sparsity value of 0.1. We observe that visual and motor cortices show a higher correspondence, which is highly myelinated. (**e**) Unimodal cortices [the superior temporal gyrus, visual (peristriate, mid-temporal, and inferior or temporal areas), and somatosensory cortex] show stronger structure-function correspondence than transmodal cortices. (**f**) A null model (permutation test, with 10000 repetitions) is performed to determine which module is statistically significant. The significant module is colored by *bright red*.

### Functional network

Two types of functional networks are constructed. For a *static functional network*, we utilize the Gordon parcellation with 333 regions as nodes (Gordon et al., 2016), and the Pearson correlation coefficients between paired nodal median time series are calculated as edges, obtaining an asymmetric connectivity matrix for each participant. For *dynamic functional networks*, the sliding window method is used to construct the functional brain network within each window; here, we use the time window with a length of 200 TRs and stepped with 17 TRs, generating 59 dynamic functional networks (***SI Appendix***, ***Materials and Methods***).

### Correspondence of microstructural covariance and mean functional network

To estimate the correspondence of microstructural covariance and mean functional network, a multilinear regression model reported by previous study (Vazquez-Rodriguez et al., 2019) is employed to uncover the node-level microstructural covariance-mean functional correspondence (**Fig. 1b**). The dependent variable of a node *i* is its resting-state FC between node *i* and all the remaining nodes in the network (*j* **≠** *i*). For the same node *i*, three properties derived from binary microstructure network, including the path length, Euclidean distance, and communicability, are used as the predictor variables. The model parameters (regression coefficients for each of the 3 predictors) are then estimated via ordinary least squares (**Fig. 1b**). For each node *i*, goodness of fit is used to represent the structure-function correspondence, which is quantified by the adjusted *R^2^* (***SI Appendix***, ***Materials and Methods***).

The correspondence of microstructural covariance and mean functional network is highly variable across the neocortex. The median value *R^2^* is 0.209 (range from 0 to 0.553), roughly concordant with previous reports using the diffusion MRI (dMRI) data to predict the whole-network FC (Goni et al., 2014) and similar pattern with the nodal structure-function correspondence (Vazquez-Rodriguez et al., 2019). However, our results show that the *R^2^* values vary extensively, indicating that for the correspondence of microstructural covariance and mean functional network, some regions own a strong correspondence, while for others, there is little evidence of any such correspondence. Our results are robust across different sparsity values (10%, 20%, 30%, and 40%) of the microstructural covariance network (**Fig. 1c**). And the main result with a sparsity value of 0.1 is shown in **Fig. 1d**; the occipital and paracentral cortices show relatively high structure-function correspondence, while the temporal cortices and some cortices within the default mode network (Raichle et al., 2001) have the least structure-function correspondence. Statistical analysis reveals that the unimodal cortices [auditory (superior temporal gyrus), visual (peristriate, mid-temporal, and inferior or temporal areas), and somatosensory cortex] show stronger microstructure-function correspondence than transmodal cortices (**Fig. 1e**, *P* < 10^-10^). To determine which module or sub-class (Gordon atlas with 13 modules, **Fig. 1c**) show non-trivial microstructure-function correspondence than null model, we use the permutation test, modules’ labels with a value from 1 to 13, are randomly shuffled (10,000 repetitions) for each brain region, we then express each sub-network mean *R^2^* as a z-score relative to this null distribution. Visual and SMhand sub-networks show significantly higher microstructure-function correspondence than random, and FrontoParietal, Auditory, and CinguloOperc sub-networks show the least microstructure-function correspondence (**Fig. 1f**).

However, these group-level structure-function interactions cannot detect individual brain-cognition relations. An individual analysis is needed. The following sections will focus on the individual microstructure-function relationships, especially on the higher-order microstructure-function correspondence.

### Myelin map and network topology

Before the exploration of the individual microstructure-function interactions, the myelin map and network topology for microstructural and functional network are introduced. We utilize the T1w/T2w ratio to depict cortical myelin content (Glasser & Van Essen, 2011). The average myelin map of each region across 198 participants demonstrates that the heavily myelinated areas are mainly in motor-somatosensory and visual cortex. By contrast, the lightly myelinated areas are classical multimodal association cortices, including prefrontal and superior parietal cortices (**Fig. 2a**). Furthermore, the average myelin content of each region is positively correlated with the correspondence of microstructural covariance and the mean functional network (r = 0.222, *P =* 4.530×10^-5^), indicating that the highly myelinated regions own stronger structure-function interactions. The 198 participants in the HCP data includes three age groups: 22-25 (M:35, F:11), 26-30 (M:50, F:43), 31-35 (M:23, F:36) years old. The mean myelin content across all regions shows an ascending trend across age groups, for the 22-25 age group is 0.898±0.063, the 26-30 age group is 0.935±0.075, and the 31-35 age group is 0.967±0.081 (**Fig. 2b**). One-way analysis of variance (ANOVA) shows a significant effect of the age group on myelin content (F _(2, 195)_ = 11.083, *P =* 2.761×10^-5^). And the female participants exhibit higher mean myelin content than the male participants (delta = 0.084, *P =* 3.603×10^-16^). See **Fig. 2c** for details.

**Figure 2.**
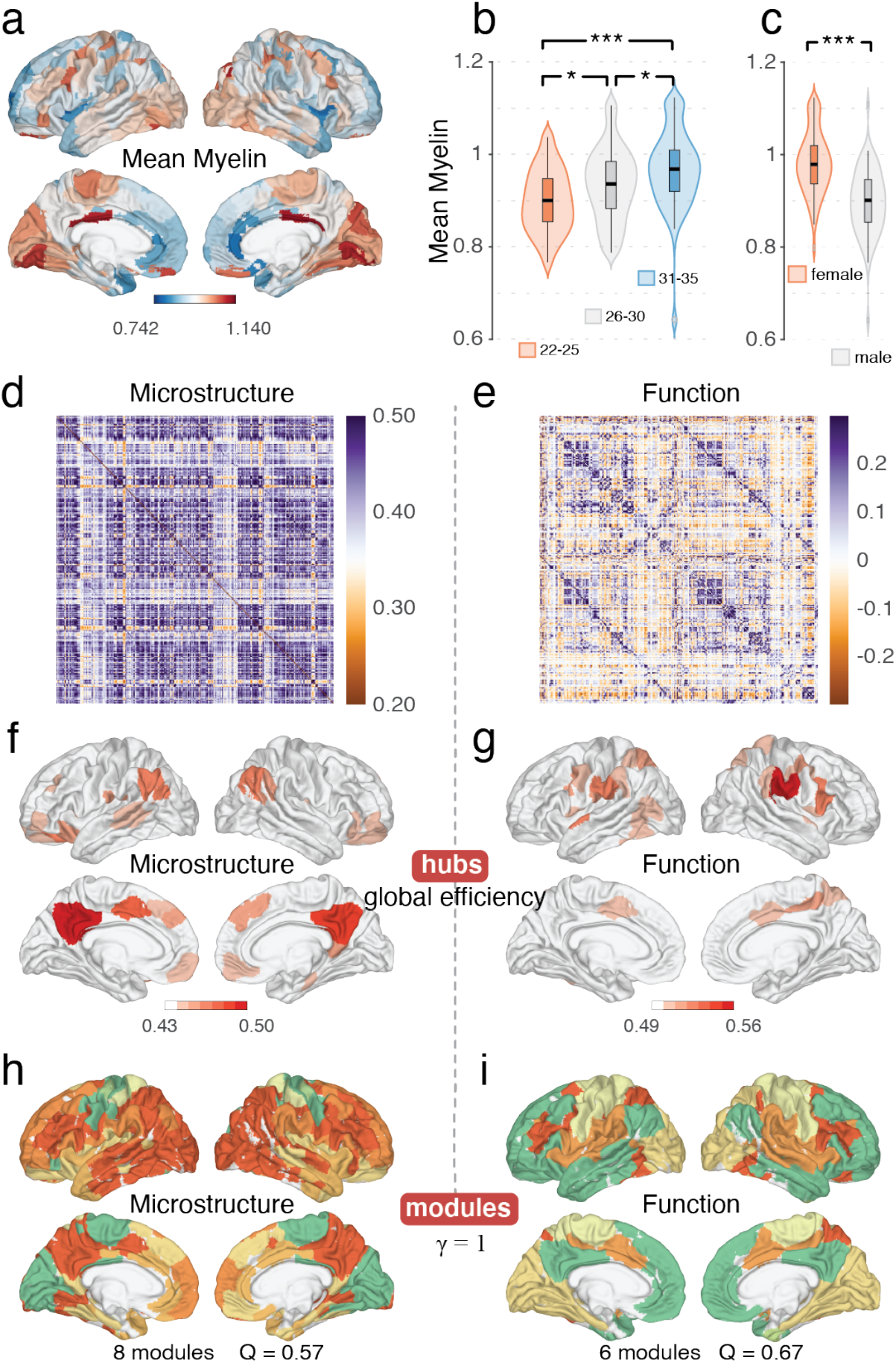
Average myelin map and network topology. (**a**) Average myelin content across all participants. (**b**) Average myelin content across three age groups. An obvious ascending trend across age groups is shown. (**c**) Average myelin content across gender. The female group shows significantly higher myelin content than the male group. (**d-e**) Mean microstructural and functional connection matrix across 198 participants. (**f-g**) Hubs of the microstructural and functional network. We term the top 10% node with a higher mean global efficiency value as hubs. (**h-i**) Modular organization of the microstructural and functional network, we observe eight modules and six modules for microstructural and functional network, respectively. *, *P* < 0.05; ***, *P* < 0.001.

The mean connectivity matrix across all participants is shown in **Fig. 2d-e** for microstructural and functional networks, respectively. We apply a sparsity threshold value of 0.1 (retaining the strongest connections in the top 10%), reported in previous studies (Grydeland et al., 2019; Lariviere et al., 2020), to convert the weighted networks to binary networks for subsequent analysis. Subsequently, we calculate four nodal metrics (clustering coefficient, local efficiency, degree centrality, and global efficiency) for each network (Rubinov & Sporns, 2010). The nodes with top 10% mean nodal global efficiency values across 198 participants are termed as hubs (Liu et al., 2017). Spatial overlapping between microstructural and functional hubs, is minimum, with a dice coefficient of 0.061 (**Fig. 2f-g)**. The community detection (Louvain algorithm) is performed on the mean microstructural and functional brain networks across all participants, with gamma (γ) = 1 (***SI Appendix***, ***Materials and Methods***). The modular similarity of the microstructural and functional network is measured by normalized mutual information (NMI), with an NMI value of 0.223 (**Fig. 2h-i)**. Besides, the small-world propensity (SWP, φ) is calculated (Muldoon et al., 2016); networks are considered small-world if they have SWP 0.4 < *φ* ≤ 1 (Bassett & Bullmore, 2017). The mean and standard deviation of SWP for microstructural network is 0.588±0.030, and for functional network is 0.691±0.031. Together, these findings indicate that both microstructure and functional networks have small-world propensity, hubs, and modular organizations, which present us with the natural question how the interactions between these two types of brain networks are, especially at higher-order level.

### Higher-order framework

Describe a network’s wiring diagram from the lower-order level alone may not capture its complex topology. Despite the same nodal degree and number of edges, two networks may have entirely different connections. Accordingly, there are four graphs with six nodes, illustrated by six different colors in **Fig. 3a**; G1 and G2 have the same degree centrality but have different topologies, while G3 and G4 have different degree centrality but are isomorphic graphs.

**Figure 3.**
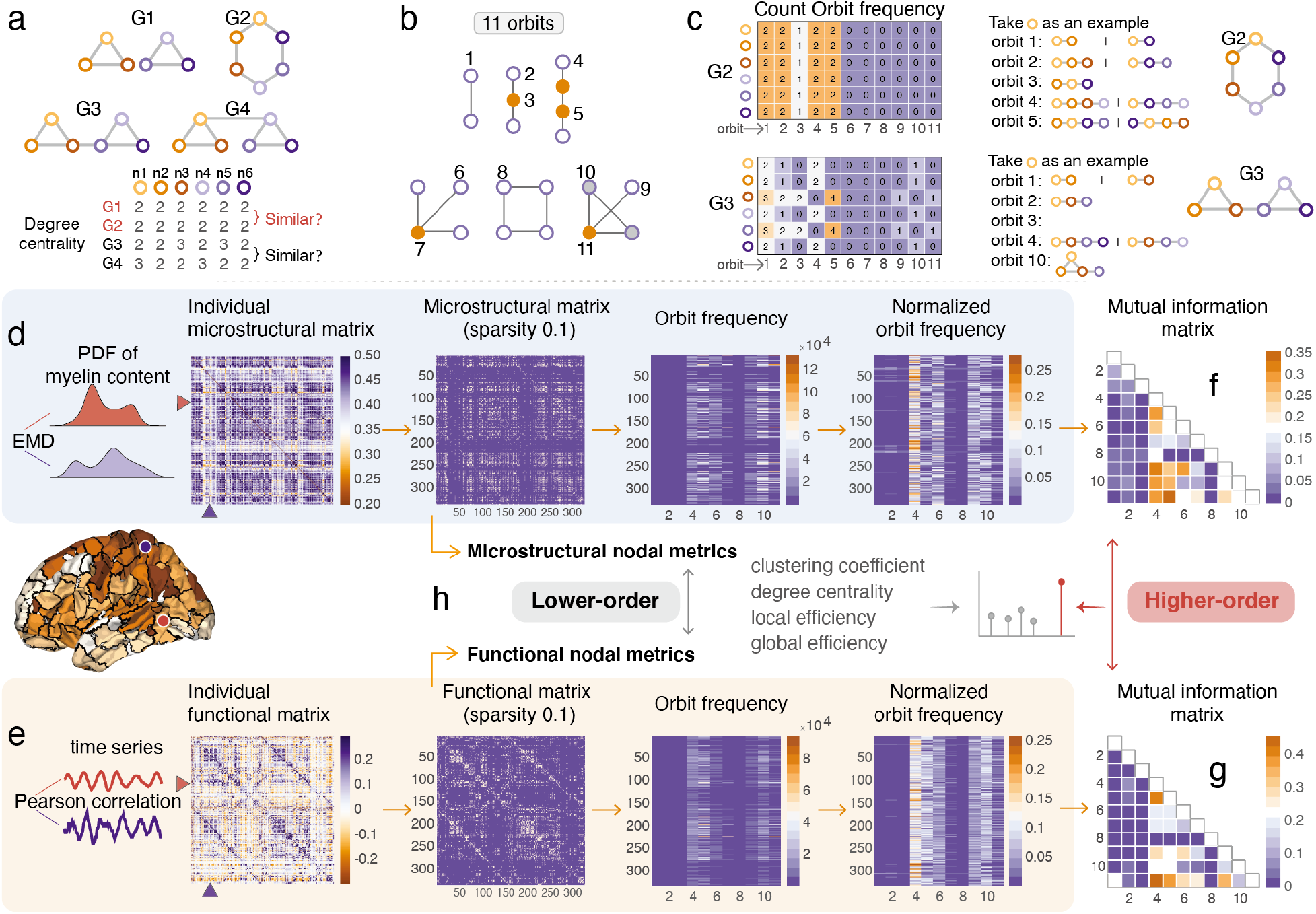
Lower- and higher-order frameworks. (**a**) Schematic representation of four simple networks with six nodes represented by different colors, *G1* and *G2* have the same number of nodes and edges, *G1* and *G2* have identical nodal degree centrality but exhibit different topological structures, *G1* consists of two triangles and *G2* is a connected circle. *G3* and *G4* own identical topological structures; they both consist of two triangles and one path, while their nodal degree centrality is different. Thus, the lower-order depiction cannot capture the whole fact. (**b**) 11 non-redundant orbits from 2-and 4-node graphlet, each orbit is derived on their unique position within a graphlet. We used 11 non-redundant orbits (Yaveroglu et al., 2014) to depict the network’s higher-order property. (**c**) Take the *G2* and *G3* as an example; we give a toy example to depict how to count the frequency of 11 orbits for each node. (**d**) Construct the individual microstructural network based on the similarity of probability distribution function (PDF) of any paired regions’ myelin content, then thresholding the network with sparsity value 10%, and calculating the orbit frequency matrix (333×11) for each network and normalizing orbit frequency matrix using logarithm (10-base) transformation. (**e**) Construct the individual functional network by calculating the Pearson correlation between the median time series of any paired brain regions, same with the structural network, we obtain the normalized orbit frequency matrix. (**f-g**) Mutual information (MI) matrix is obtained by calculating the similarity between the 11 orbits. We term the Pearson correlation between the MI matrix of the myelin-based microstructural and functional network as their higher-order relationship or interaction. (**h**) Lower-order framework, we term the Pearson correlation between the microstructural and functional brain network’s nodal metrics as the lower-order similarity.

The similarity between the two networks can be simply defined as the correlation of nodal features or edge’s features at the lower-order level. Beyond the node and edge, subgraphs, which are the essential building of a network, can be used to explore the network’s higher-order patterns. To obtain the higher-order interactions, we employed 11 non-redundant orbits (the position of nodes inside each subgraph) reported by a previous study (Yaveroglu et al., 2014), derived from 2- to 4-node subgraphs. The 11 non-redundant orbits are used here because they are most efficient in computing and perform well in clustering networks with the same topological properties (Yaveroglu et al., 2014), see **Fig. 3b.** We then give a toy example of counting the frequency (number of times a node touches a specific orbit) of 11 orbits for each node in a graph (**Fig. 3c**).

To measure the relationships of microstructural and functional brain networks at the macroscale individual level, 198 participants are used for the subsequent connectome and statistical analysis. Specifically, for the microstructural brain networks, edges are determined by calculating the earth mover’s distance (EMD) (Ruttenberg & Singh, 2011) between the probability distribution function (PDF) of nodal myelin content, based on previous studies (Leming et al., 2021; Wang et al., 2016), see **Fig. 3d**. For functional networks, edges are defined by estimating the statistical similarity between nodal time series (**Fig. 3e**), see ***SI Appendix***, ***Materials and Methods*** for details. Here, we compute the frequency of 11 non-redundant higher-order features mentioned above for each node in each microstructural and functional brain network, then the interrelationships between these 11 features are estimated by mutual information, which can describe the linear or non-linear relationship between variables, yielding an 11-by-11 mutual information matrix (**Fig. 3f-g**). The Pearson correlation between the mutual information matrices of the microstructural and functional brain networks is termed as the higher-order microstructure-function relationship or interaction. Besides, the lower-order similarity, defined as the Pearson correlation of the nodal feature of each type of network, is compared as the baseline (**Fig. 3h**).

### Global (network-level) higher-order microstructure-function relationship

We apply the higher-order framework mentioned above, to explore the higher-order microstructure-function relationship. Unless otherwise stated, the thresholded binary networks with a sparsity of 10% is used for the subsequent statistical analysis. Besides, the lower-order relationship, described by Pearson correlation of their nodal features, is also performed as a comparison result of higher-order level; two types of null random networks, preserving the microstructural and functional network’s degree distribution, respectively, are considered as the compared baseline.

#### Enhanced higher-order interactions

Higher-order framework revealing the microstructure-function higher-order relationship is 0.860±0.042 (mean value ± standard deviation), which is higher than the lower-order similarity (all *P-values* < 10^-9^, *FDR correction*) (**Fig. 4a**). Notably, at the lower-order level, microstructural and functional networks’ similarities are lower than their relationships with corresponding random networks (all *P-values* < 10^-9^, *FDR correction*). It is surprising because both the microstructural and functional networks are somehow derived from the properties of the brain and are different from random networks and may exhibit a high correlation. At the higher-order level, the microstructural and functional networks show a higher correlation than their corresponding random networks (**Fig. 4a**), implying that the higher-order method may better seize the network’s global attribute information or hidden pattern.

**Figure 4.**
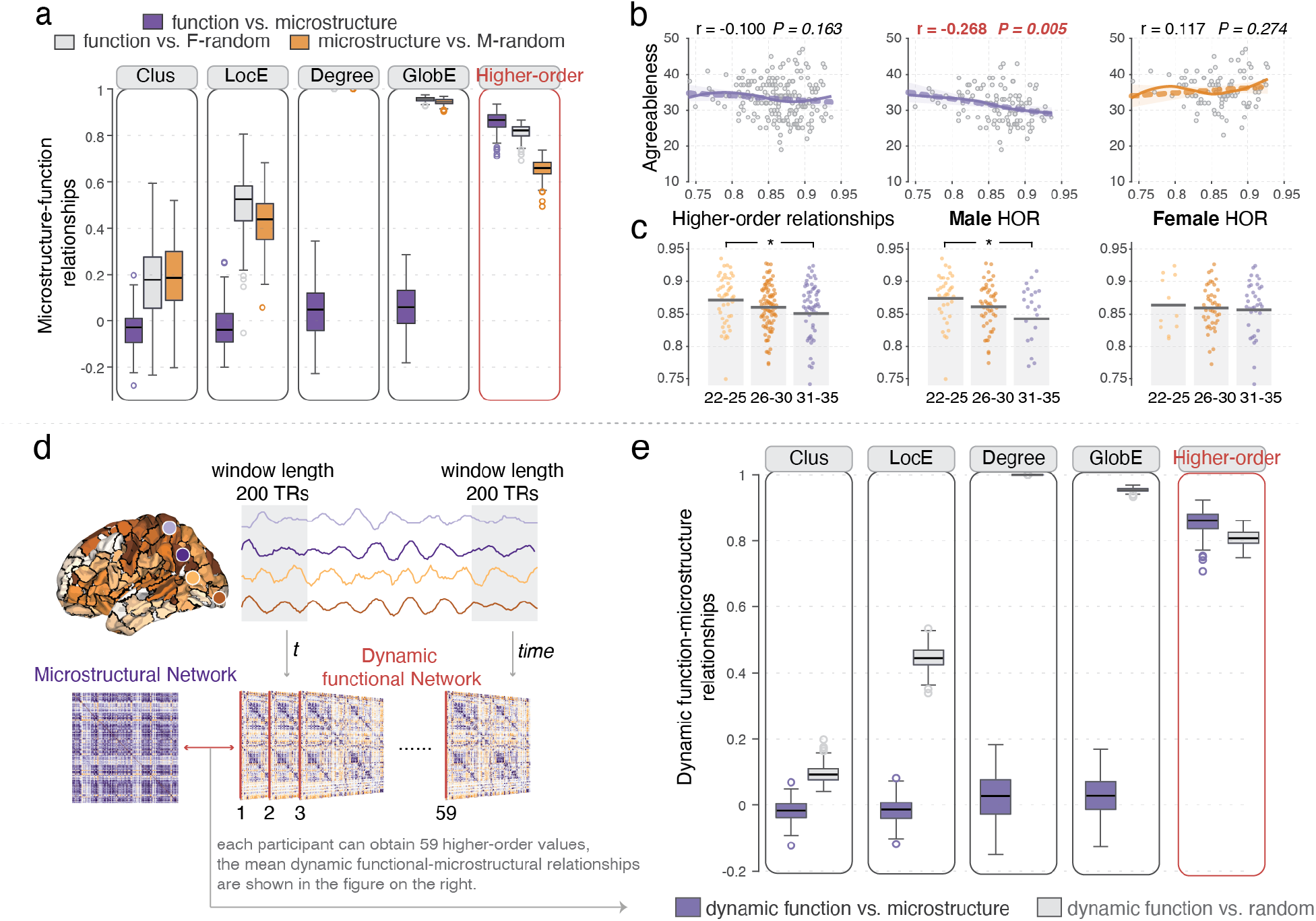
Global (network-level) higher-order microstructure-function relationship. (**a**) Lower- and higher-order relationships between the microstructural and functional network. Clus, clustering coefficient; LocE, local efficiency; Degree, degree centrality; GlobE, global efficiency. (**b**) Association between brain and cognition. Correlation between individual higher-order relationships and agreeableness for male and female participants. (**c**) Changes of higher-order relationships across three age groups. *, *P* < 0.05. (**d**) One-to-many (microstructure-functions) relationships. We use the sliding window to construct 59 dynamic functional networks for each participant and calculate the relationship between each participant’s microstructural and 59 dynamic functional brain networks. (**e**) The detailed results of lower- and higher-order relationships between the microstructural and 59 dynamic functional networks across participants, indicating the stronger higher-order interaction can be largely maintained across the different dynamic status.

#### Link cognition with higher-order interactions

Previous studies have shown that gender plays a crucial role in the personality-brain relationship (Nostro et al., 2017). To further evaluate the high-order relationship’ physiological significance, we use the NEO Five-Factor Inventory (NEO FFI) (Costa & McCrae, 1989), which includes five dimensions: openness, conscientiousness, extraversion, agreeableness, and neuroticism, to correlate with the individual higher-order relationship across the whole participants and separately for males and females. We observe the higher-order relationship is significantly negatively correlated with agreeableness across the male participant (r = -0.268, *P =* 0.005) rather than female participants, see **Fig. 4b**, indicating that the higher-order relationship correlated with personality is highly dependent on gender. In this study, we use the unrestricted HCP dataset, which includes three age groups: 22-25 (M:35, F:11), 26-30 (M:50, F:43), 31-35 (M:23, F:36) years old. The ANOVA test is performed to verify whether the microstructure-function higher-order relationships were significantly different across the three groups, and a decreasing trend is detected; the mean value of higher-order relationships is 0.872 for the 22-25 group, 0.861 for the 26-30 group, and 0.851 for the 31-35 group (**Fig. 4c**). The ANOVA analysis shows there is a significant difference in higher-order relationships among the three groups (F _(2,195)_ = 3.197, *P* = 0.043) and the *post-hoc* indicates that the 22-25 group exhibits a stronger higher-order relation than the 31-35 group (delta = 0.021, *P*_tukey_ = 0.031). The male participants mainly drive this effect. We do not observe significant results for the female participants. Together, these findings once again demonstrate the pivotal role of age and gender in higher-order relationships.

#### Higher-order interactions of microstructure-dynamic functional networks

To investigate the one-to-many (microstructure-functions) relationships, we use the sliding window method (window length is 200 TRs and step length is 17 TRs) to construct 59 dynamic functional brain networks (Preti et al., 2017) and evaluate their relationships with the corresponding static myelin-based microstructural network for each participant (**Fig. 4d**). Similar patterns are obtained compared to the one-to-one (microstructure-function) analysis; the mean higher-order relationship between the static microstructural and 59 dynamic functional networks of each participant is higher than their relationships with the random networks (**Fig. 4e**). It reveals that even in different states, the strength of structural and functional interactions is higher than that in random networks, which may be a principle of how the brain works.

### Nodal (node-level) higher-order microstructure-function relationship

The nodal (regional) higher-order relationships are defined by the cosine similarity between the frequency of 11 orbits of each node in functional and microstructural network (**Fig. 5a**). To determine whether the higher-order microstructure-function relationship depends on the structural or functional nodal centrality. We perform the correlation analysis between the nodal higher-order relationship and nodal four centrality metrics (clustering coefficient, local efficiency, degree centrality, global efficiency). Significant positively correlations are detected (all *P-values* < 0.005, *FDR correction*), demonstrating that nodal centrality measures are associated with the observed nodal higher-order microstructure-function patterns (***SI Appendix***, **Fig. S1**). We find that the nodal higher-order relationships are stronger in default mode, visual, and frontoparietal regions and weaker in the ParietalOccip and somatomotor cortex (**Fig. 5b**, *permuted P < 0.01*). And the map of the nodal higher-order microstructure-function relationship (**Fig. 5a**) is negatively correlated with the spatial distribution map of structure covariance-function correspondence **(Fig. 1d**), as shown in **Fig. 5c.**

**Figure 5.**
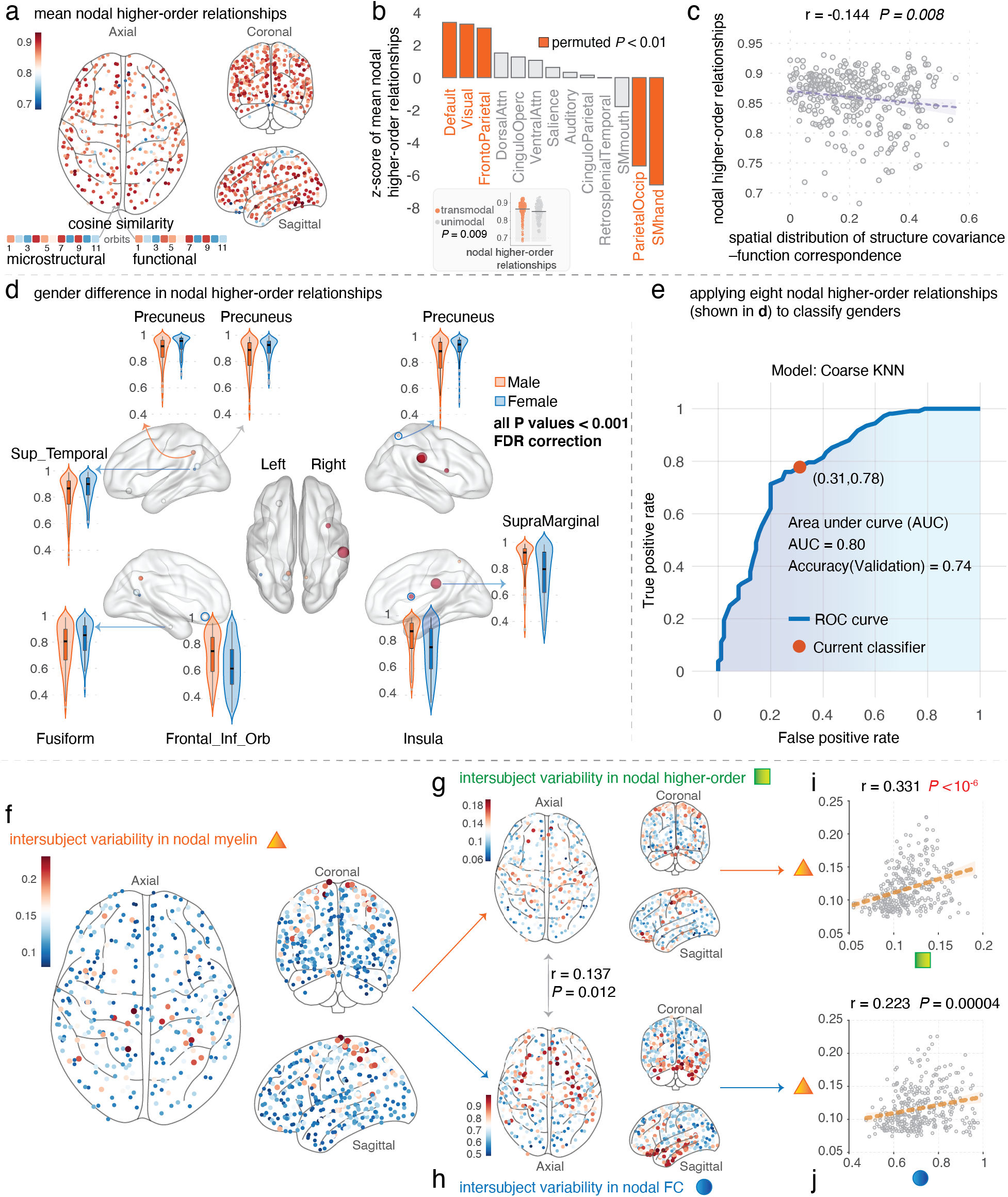
Nodal (node-level) higher-order microstructure-function relationship. (**a**) Nodal (regional) higher-order relationships were defined by the cosine similarity between the frequency of 11 orbits of each node in the functional and microstructural network. (**b**) Z-score of nodal higher-order relationships. (**c**) Nodal higher-order microstructure-function relationship is negatively correlated with the spatial distribution map of structure covariance-function correspondence. (**d**) Gender difference in nodal higher-order relationships. (**e**) The performance of classification in nodal higher-order relationships. *Individual variability* in nodal myelin (**f**), higher-order relationships (**g**), and functional connectivity-FC (**h**). The correlation between nodal microstructural variability and the variability in nodal higher-order interactions (**i**) and the variability in nodal FC (**j**).

#### Gender difference

Due to the different needs and purposes of male and female social behavior, we might expect these behaviors to lead to specific neurocognitive adaptations (Kiesow et al., 2020). Here, we probe the gender difference in nodal higher-order microstructure-function interactions, some of the left (precuneus, fusiform, superior temporal gyrus, inferior frontal gyrus-orbital part) and the right (precuneus, insula, and supramarginal gyrus) regions show significant differences in nodal higher-order microstructure-function interactions (all *P-values* < 0.001, *FDR correction*) (**Fig. 5d**) and we can use these nodal higher-order interactions to classify gender group with an accuracy of 0.74 and AUC of 0.80 by *coarse KNN* (K nearest neighbor) model, executed in MATLAB 2021a classification learner app with parameters: number of neighbors: 100, distance metric: Euclidean, distance weight: equal, and standardize data: true (**Fig. 5e**).

#### Individual variability

A previous study shows that FC variability is related to anatomical variability (Mueller et al., 2013), which may explain individual differences in cognition and behavior. Here, we inspect the intersubject variability in nodal higher-order interactions and nodal FC and link them with the microstructural variability to identify whether their intersubject variability can be partially explained by microstructural variability. We term the *standard deviation* of each regional myelin content across all 198 participants as the nodal microstructural variability (**Fig. 5f**). Individual microstructural variability is larger in the motor cortex and less within the default mode network. Furthermore, we define the *standard deviation* of nodal higher-order microstructure-function relationships across all 198 participants as the variability of nodal higher-order interactions, intersubject variability in nodal higher-order microstructure-function relationship is stronger in the somatomotor cortex (**Fig. 5g**). Subsequently, for the FC, we calculate each node’s stability, which is indicated by the *intraclass correlation coefficient* (ICC) of this node with all other nodes’ FC across all participants (332 × 198). We define the variability in nodal FC as 1–ICC (**Fig. 5h**), which is greater in the transmodal association cortex, including the prefrontal and temporal lobe, and lower in the unimodal visual and motor cortices. Nodal microstructural variability (**Fig. 5f**) shows a moderate correlation with the variability in nodal higher-order microstructure-function interactions (r = 0.331, *P* < 10^-6^), see **Fig. 5i** for details, and significant correlation with the variability in nodal FC (r = 0.223, *P* = 0.00004), *FDR correction*, see **Fig. 5j** for details.

### Local (circuit-level) higher-order microstructure-function relationship

Higher-order microstructure-function relationships are stronger compared with their corresponding random networks for the whole brain. To check whether this property still is held for local circuits, we use random sampling (100 iterations) to extract subnetworks from the whole microstructural and functional brain networks. The sampling number of nodes decreased from 240 to 30 (interval of 5), yielding 43 steps. We obtain 43×100 subnetworks for each participant’s microstructural and functional networks, respectively. Then, we calculate the microstructure-function higher-order interaction of those subnetworks for each step for each participant (see **Fig. 6a-c**, results of networks with 240, 120, 60, 30 nodes are shown). We find that the subnetworks’ microstructure-function higher-order relationships are significantly higher than their relationships with random networks, regardless of the nodal resolution (from 240 to 30 nodes, see ***SI Appendix*, Fig. S2**). These findings confirm those observed in whole brain networks and indicate that both global and local circuits exhibit stronger microstructure-function higher-order interactions than random networks, which maybe a vital principle for human brain.

**Figure 6.**
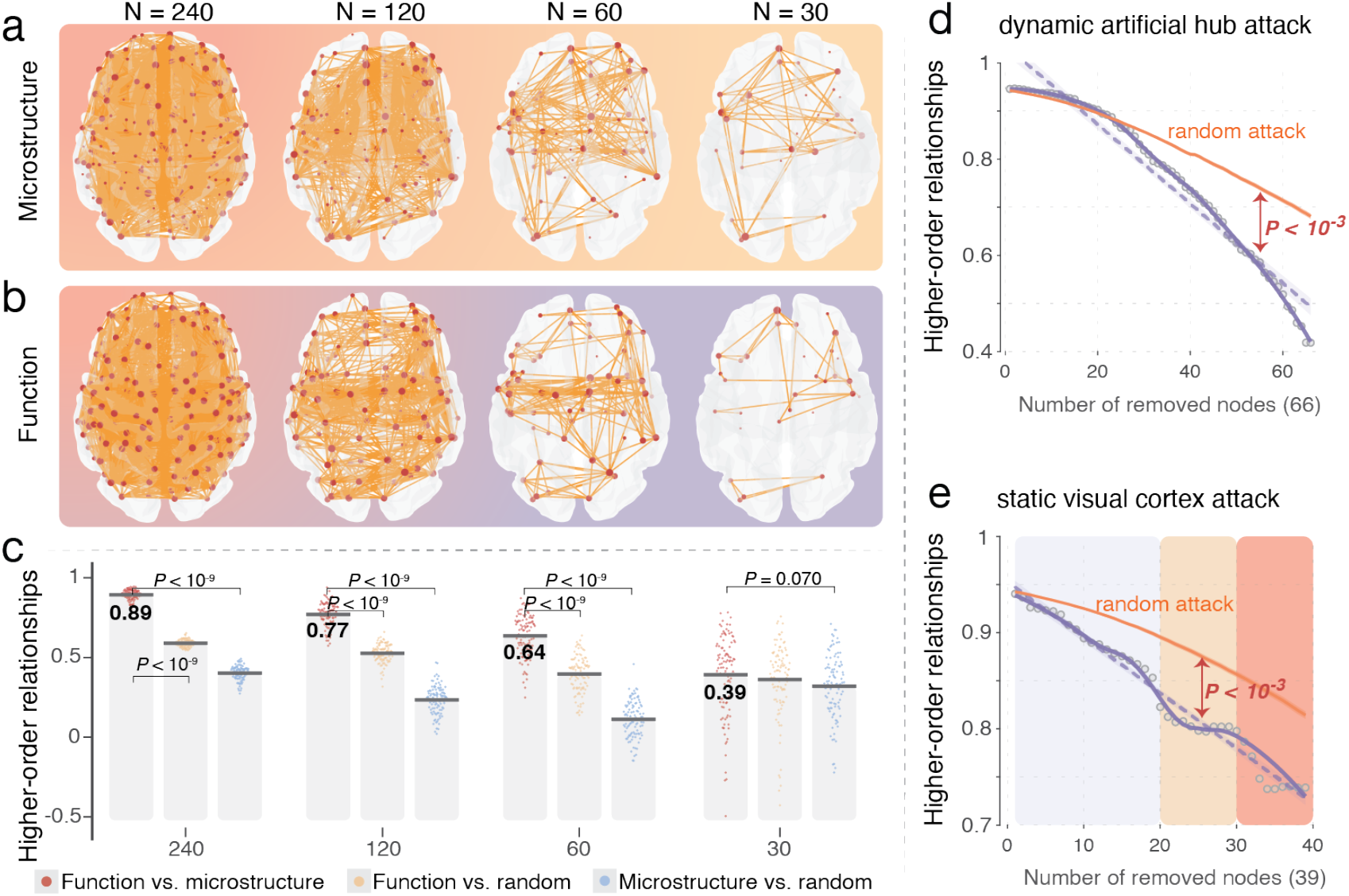
*Local (circuit-level)* higher-order microstructure-function relationship and *resilience analysis*. (a) We randomly sample 100 times with different resolutions (network size) to extract the microstructural circuits (subnetworks); Here, the schematic shows the results for only 240,120, 60, and 30 nodes for one-time sampling. (**b**) The node index is consistent between the microstructural and functional network; the functional circuits (subnetworks) are also shown for only 240,120, 60, and 30 nodes. (**c**) After random sampling, we calculate the higher-order relationships between the microstructure-function subnetwork and their relationships with the random networks. (**d**) We performed targeted attacks for vital nodes with higher global efficiency in the mean microstructural network one-by-one until we removed 20% nodes of the network and recalculated the higher-order interactions with the mean functional network, showing hub attacked can significantly disrupt the higher-order interaction than random attacks. (**e**) We also performed the static visual cortex attack, removed the nodes one by one, in descending order for global efficiency, only in the microstructural network. Roughly, three stages of change can be observed. The dashed line indicates linear (least squares) fit; the solid line indicates non-parametric regression [LOESS (locally estimated scatterplot smoothing)] fit.

### Resilience of structure-function higher-order relationships

Previous studies showed that hub lesion/attack (an essential indicator of brain disorders) caused the most extensive brain network organization disturbances (Aerts et al., 2016; Morone & Makse, 2015). Here, we perform two kinds of attacks: *dynamic artificial hubs attack* and *static visual cortices attack* to investigate the resilience of microstructure-function higher-order interactions. We use the mean microstructural and functional network, thresholding to only keep the top 10% edges in the full connection matrix; we do not modify the functional network and the lesion is just performed on the microstructural network. For the *dynamic artificial hub attack*, we first remove the node with the highest global efficiency, setting all connections to that node to 0, then calculating the higher-order interaction between the lesioned microstructural network and the functional network. Our next step is to calculate the global efficiency of every node in the lesioned microstructural network and remove the node with the highest global efficiency.

Then recalculate the higher-order interactions between the nodes until we have deleted 20% nodes of the microstructural network. The results show that *dynamic artificial hub attack* significantly disrupts the microstructure-function higher-order relationships compared with random attacks (1000 iterations with random attacks) (**Fig. 6d**). The visual cortex contains a high myelin content, but some demyelinating diseases, such as multiple sclerosis exhibit abnormalities in the visual areas. To determine whether the changes of visual microstructural content will affect the higher-order microstructure-function interactions. Here, we perform a static attack on the visual cortices; the *static visual cortex attack* is carried out by ranking the nodal global efficiency within the visual cortex in descending order, then deleting them one by one only in the microstructural network, recalculating the microstructure-function higher-order relationships for each deleting, until all 39 visual regions have been deleted. Results show that *the static visual cortex attack* significantly disrupts the microstructure-function higher-order relationships than the random attack (**Fig. 6e**). Three stages are obtained: early (1-20 lesioned nodes); middle (20-30 lesioned nodes); and late (30-39 lesioned nodes), which better matches the clinical stage of demyelinating disease (**Fig. 6e**). Taking multiple sclerosis as an example, which is the most common demyelinating disease of the central nervous system, broadly comprises three stages: (1) a pre-clinical stage; (2) a relapsing-remitting (RRMS) clinical stage; and (3) a progressive clinical-stage during which neurologic dysfunction progressively worsens (Baecher-Allan et al., 2018). This simple simulation attack in the visual cortex can disrupt higher-order structure-function interactions, which may help understand some demyelinating diseases from a structure-function interactions perspective. Overall, these two simple models can significantly disrupt the higher-order interactions than random attacks (all *P-values* < 0.001, *permutation test* for comparing two curves). Jointly, the higher-order framework demonstrates potential application in monitoring disease progression. Further insights will be gained from studies that utilize the higher-order framework with real clinical data rather than simulated data, such as multiple sclerosis, traumatic brain injury, and Alzheimer’s disease.

### Additional analysis

We also performed two additional analyses to better understand the higher-order framework: 1) Apply the higher-order method to network classification to test the sensitivity of this method.

Results prove the higher-order framework performs efficiently in classifying networks. 2) Explore the relations between nodal metrics and 11 orbits to gain a deeper understanding of the higher-order features. See ***SI Appendix, Additional analysis,* Fig. S3**.

### Robustness

The robustness of our main results is examined against several factors, including (I) effect of sparsity thresholding; (II) effect of sample size; (III) split-half reliability for nodal higher-order interactions; (IV) higher-order relationships on other different types of networks; (V) effect of different embedding methods; (VI) reliability of constructing the individual microstructural network. Taken together, the results of our study are largely reliable and robust; see ***SI Appendix, Robustness*, Figs. S4-S10** for details.

## DISCUSSION

This work systematically characterized the relationship of myelin microstructure and function from the population-level to individual-level, using both the lower-order and higher-order frameworks. In particular, the higher-order microstructure-function relationship is significantly higher than their relationships in random networks. Interestingly, this stronger higher-order microstructure-function relationship can be maintained from whole-brain to local circuits even when the circuit size is decreased, which maybe an important principle of human brain.

Furthermore, higher-order microstructure-function relationship is not uniform across the brain, showing differences between genders, and individual variability is related to inter-subject variability. In addition, a targeted attack may disrupt the higher-order microstructure-function relationship. These results provide new higher-order insights into understanding the interactions between brain microstructure and function (Holler et al., 2021; Suarez et al., 2020), linking these interactions to cognition and behavior, and driving the development of brain-like computations and applications into discovering relationships among different complex systems in other disciplines.

### Microstructure covariance–function correspondence

The myelin content has increased among all age groups from 22-25 to 31-35 years, consistent with recent studies (Grydeland et al., 2019). During development, myelination occurs within the cortex and reaches a point of stability at 30 years of age, following which it declines at 60 years of age (Grydeland et al., 2019; Melie-Garcia et al., 2018). Moreover, our results suggest that node-level structure-function correspondence of microstructural covariance network and mean functional network is not aligned uniformly across the brain. Higher structure-function correspondence in unimodal cortices (motor and visual cortex), lower structure-function correspondence in some transmodal cortices (frontal-parietal and default mode regions), which is comparable with previous study (Vazquez-Rodriguez et al., 2019). There is a consensus that individual areas have distinctive “fingerprints” (Finn et al., 2015) and different modules (Avena-Koenigsberger et al., 2017). The somatosensory, motor, and early visual cortex have the highest myelin content, while the association / multisensory cortex has the lowest (Fukutomi et al., 2018; Glasser & Van Essen, 2011; Hunt et al., 2016). Research suggests that cortical myelin possibly inhibits plasticity. For example, early sensory areas may require less plasticity and more myelin, while higher-order areas may have less myelination and therefore be more plastic (Glasser et al., 2014). It appears that myelin acts to increase processing speed and inhibit plasticity in primary sensory areas (Glasser et al., 2014). In that way, it sheds light on the mechanism by which cortical microstructure supports functional networks.

### Individual higher-order microstructure–function relationship

As opposed to previous studies based on group-level (Hunt et al., 2016; Ma & Zhang, 2017; Melie-Garcia et al., 2018), we constructed individual-level myelin-based microstructural networks for direct comparison between individuals. The results demonstrate that these microstructural networks are characterized by small-world propensity (Bassett & Bullmore, 2017), hubs, and modular organization, which provide a foundation for exploring the structure-function relationship. To deepen the systematic understanding of the complex structure-function relationships, we examine the one-to-one (microstructure-function) and one-to-many (microstructure-functions) patterns, from the lower-order to the higher-order level. The lower-order relationships may reflect the functional brain network’s flexibility due to copy with different states (Park & Friston, 2013), and the higher-order may capture the hidden pattern of networks. Results show that, compared with microstructure-random and function-random relationships, the microstructure-function (s) relationship is smaller at the lower-order level. This result is consistent with previous structure-function research at the node and edge level (Di et al., 2017; Hunt et al., 2016).

However, it becomes larger at the higher-order level. Furthermore, we find the global higher-order relationships are associated with individual personality scores and descended with aging, and male participants primarily drive the result. Consistent with the previous study (Nostro et al., 2017), it highlights the brain structure–personality relationships are highly dependent on gender.

For nodal higher-order microstructure-function relationships, default mode network regions show stronger higher-order microstructure-function relationships, while motor cortices show weaker higher-order microstructure-function relationships. The transmodal regions show stronger nodal higher-order microstructure-function relationships than unimodal regions, which has an opposite pattern to node-level microstructural covariance-function correspondence, especially for the default mode regions (Raichle et al., 2001), which have weaker microstructural covariance-function correspondence while having stronger nodal higher-order microstructure-function interactions, providing a new perspective and tool to understand the default mode network (Raichle et al., 2001) and the structure-function interactions of association cortex (Sydnor et al., 2021). The regions like precuneus, fusiform, and insula, observed in previous sex differentiation study with large participants (Kiesow et al., 2020), show significant gender effect in nodal higher-order microstructure-function relationships. We can use these regions to distinguish the male from the female with good accuracy, highlight the potential application in different groups classification and some clinical monition for brain disorder participants.

Additionally, a modest but significant relationship was observed between sulcal depth variability and functional variability (Mueller et al., 2013) in a previous study (r = 0.30, *P* < 0.0001). We observe very similar results in the correlation between myelin variability and higher-order microstructure-function variability, indicating that individual structure differences may affect the structure-function interactions.

For structure-function higher-order interactions of local circuits. The microstructure-function higher-order relationships can be preserved even with a gradually decreasing scale. In this study, both whole-brain networks and local circuits exhibit stronger higher-order relationships than their relationships with random networks. The important principle, greater structure-function interactions from whole-brain to local circuits, may help design new and efficient brain-inspired intelligence and further understanding the role of structure-function higher-order interactions of local circuits in human aging, cognitive activities, and personality.

### Potential applications and targeted attack

The higher-order framework can classify different types of networks with higher accuracy than the lower-order one, exhibiting the potential to construct a network of networks, describing the relationships among different complex systems. And the targeted attack (Aerts et al., 2016; Crossley et al., 2014) can disrupt the higher-order microstructure-function relationships and the visual cortices attack yields decreased higher-order relationship in a three-stage curve process, the simple simulated models are match for the evolution of demyelinating diseases such as multiple sclerosis (Baecher-Allan et al., 2018), providing us with confidence in monitoring disease progression with higher-order structure-function relationships.

### Additional considerations and perspectives

Several methodological limitations and considerations affect the present results. *First*, our estimation of myelin is based upon T1w/T2w ratio, although serving as an efficient marker of myelination, it is important to note that there is no one-to-one relation between T1w/T2w ratios [or magnetization transfer rate (MTR) or the grey/white matter contrast (GWC)] and myelin density (Glasser & Van Essen, 2011). *Second*, we only compare the resting-state functional brain network and the microstructural network; the pattern of microstructure-function relationship under the various tasks-based functional network is largely unknown. *Thirdly*, artificial attacks are made to investigate the changes of higher-order relationship, nevertheless, the models will not accurately reflect the real situations, applying the higher-order framework to trauma injury or demyelinating disease (like multiple sclerosis) participants could give us important new comprehensions. Research utilizes different myelin imaging approaches, such as MTR or GWC, to construct individual microstructure networks to inspect the microstructure-function relationship is warranted. As well, enrolling task-based data or developing data with a larger age range (e.g., 5-80 years old) will undoubtedly yield more insights. For applications, the global, nodal, or local circuits’ structure-function relationship may serve as a potential indicator of disease progression.

## CONCLUSION

These findings suggest that examining myelin microstructure-function patterns and how they relate to cognition and aging is feasible and indeed desirable. With this foundation, human brain connectome studies can move beyond the low-order (node or edge) level of inferences, to reveal the principle of structure-function interactions at the individual higher-order (subgraphs, motifs, hypergraphs, etc.) level, understand ongoing structure-function interactions with brain development or disorder, and inspire the development of brain-like intelligent computing framework. Furthermore, the method we used can also be applied to other fields (physics, economics, etc.) to measure how similar different complex systems are, and network classification, etc.

## MATERIALS AND METHODS

For details, see *Supplementary Information*.

## CONFLICT OF INTERESTS

The authors declare no competing financial interests.

## FUNDING

This work is supported by the National Natural Science Foundation of China (Nos. 61673150, 11622538), the Science Strength Promotion Program of the University of Electronic Science and Technology of China (No. Y030190261010020), and the China Scholarship Council (No. 201906070121).

## ACKNOWLEDGMENTS

We thank the investigators and staff members of the Human Connectome Project consortium for their invaluable contributions to data acquisition and sharing.

## AUTHOR CONTRIBUTIONS

Conceptualization: H.W. and L.L.; Methodology: H.W., Y.Y.L., and L.L.; Investigation: H.W. and L.L.; Visualization: H.W. and H.J.W.; Supervision: L.L.; Writing-original draft: H.W. and H.J.W.; Writing-review & editing: Y.Y.L., and L.L. All authors reviewed the manuscript.

## DATA AVAILABILITY

Human Connectome dataset: Data were provided by the Human Connectome Project, WU-Minn Consortium (Principal Investigators: David Van Essen and Kamil Ugurbil; 1U54MH091657) funded by the 16 NIH Institutes and Centers that support the NIH Blueprint for Neuroscience Research; and by the McDonnell Center for Systems Neuroscience at Washington University (https://doi.org/10.1038/nn.4361).

## CODE AVAILABILITY

The T1-weighted and T2-weighted data were processed using the MRTool (https://www.nitrc.org/projects/mrtool, version 1.4.2) implemented in the SPM12 (https://www.fil.ion.ucl.ac.uk/spm/software/spm12, version 7771). Graph metrics were implied with BCT toolbox (https://sites.google.com/site/bctnet, version 2019-03-03) and Custom-written MATLAB codes for calculating the microstructure–function higher-order relationships are freely available at https://github.com/HW-HaoWang/higher-order-interaction.

## MATERIALS AND METHODS

### MRI data

Dataset: We download unprocessed MR data of 213 participants from the Human Connectome Project (HCP) “S1200” new subjects release, the HCP (PI: David Van Essen and Kamil Ugurbil; 1U54MH091657) was funded by the 16 NIH Institutes and Centers that support the NIH Blueprint for Neuroscience Research, and by the McDonnell Center for Systems Neuroscience at Washington University (Van Essen et al., 2013), and these DICOM files were converted to NIfTI format using the dcm2nii utility (http://www.nitrc.org/projects/mricron).

Informed consent was obtained from all participants, and participants were recruited from Washington University (St. Louis, MO) and the surrounding area. We excluded eight participants who met any of the following criteria: (a) mean of framewise displacement (mFD) > 0.25 mm; (b) more than 20% of the FDs were above 0.2 mm; and (c) if any FDs were greater than 5 mm (Jenkinson, Bannister, Brady, & Smith, 2002; Parkes, Fulcher, Yucel, & Fornito, 2018). Seven participants > 36 years old were also excluded. Finally, we obtained quality-controlled resting-state fMRI (rfMRI) images, T1-weighted images, and T2-weighted images of 198 participants (108 males and 90 females, see the ***SI Appendix,* Table S1** for a complete list of participant IDs).

### Structural data

Structural scans with the following parameters were also collected: T1-weighted (0.7 mm isotropic resolution, TR = 2400 ms, TE = 2.14 ms, flip angle = 8 deg, FOV = 224 × 224 mm, acquisition time = 7min 40 sec, BW = 210 Hz/Px) and T2-weighted (0.7 mm isotropic resolution, TR = 3200 ms, TE = 565 ms, variable flip angle, FOV = 224×224 mm, acquisition time = 8 min 24 sec, BW = 744 Hz/Px) data.

#### Structural data preprocessing

The T1-weighted and T2-weighted data were processed using the MRTool (Ganzetti, Wenderoth, & Mantini, 2014) (https://www.nitrc.org/projects/mrtool, version 1.4.2) implemented in the SPM12. MRTools provided bias correction and intensity calibration on both images and then calculated the ratio between preprocessed T1w and T2w images.

Finally, we obtained the T1w/T2w ratio images in the MNI space with a resolution of 1 × 1 × 1 mm resolution.

#### Construct the microstructural covariance network

We first used the Gordon parcellation with 333 brain regions (Gordon et al., 2016) to extract the mean value of myelin (T1w/T2w ratios) content for each region of each participant, obtaining a 333 × 198 matrix, linear regression was performed on the regional mean myelin to remove the effects of age groups and gender, then the residuals of this regression was used to perform the Spearman correlation on each paired regions across the 198 participants, resulting into a 333 × 333 matrix. The framework measures the covariation of information from different brain regions across a population, known as the covariance network (He, Chen, & Evans, 2007), and thus we acquired a group-level microstructural covariance network (**Fig. 1a**).

#### Construct the individual myelin-based microstructural network

Same with the functional brain network, we used the same brain atlas (Gordon atlas with 333 regions) for myelin-based microstructural network analysis. For each participant’s myelin map (T1w/T2w ratio) map, we first extracted the myelin values for all voxels within each brain region. For each brain region, we used kernel density estimation (KDE) to estimate the probability density function (function: kde.m, https://www.mathworks.com/matlabcentral/fileexchange/14034-kernel-density-estimator) of these myelin values with 128 sampling points (Wang, Jin, Zhang, & Wang, 2016) and the bandwidth of the range interval is the minimum to the maximum of the whole brain myelin value. We standardized the probability density function by dividing its sum to produce a probability distribution function (PDF) to ensure that the PDF sum is 1. Subsequently, we calculated the Earth Mover’s Distance (EMD) (*D_E_*) between the PDFs of any paired brain regions, resulting in a 333 × 333 matrix. We converted the EMD matrix to a similarity matrix based on sigmoid function and termed the myelin-based similarity *M_sim_* for each participant as

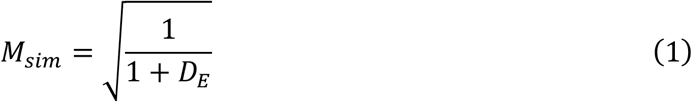

### Functional data

The rfMRI data was collected using a 32-channel head coil on a modified 3T Skyra system (MAGNETOM Skyra Siemens Healthcare). Scanning Sequence is Gradient-echo EPI, with a multi-band acceleration factor of 8, TR = 720 ms, TE = 33.1 ms, flip angle = 52 deg, FOV = 208 × 180 mm (RO × PE), matrix = 104 × 90 (RO × PE), slice thickness = 2.0 mm; 72 slices; 2.0 mm isotropic voxels, Echo spacing = 0.58 ms, BW = 2290 Hz/Px. The rfMRI data was acquired in four runs of 14 min 33 sec each, two runs were included in one session, and two were included in another session, with eyes open condition. Within each session, each run had a different phase encoding in a right-to-left (RL) direction or a left-to-right (LR) direction (REST1_LR, REST1_RL, REST2_LR, and REST2_RL). In the current study, we only used the “REST1_RL” unprocessed data.

#### Functional data preprocessing

Data were processed using SPM12 (version r7219, http://www.fil.ion.ucl.ac.uk/spm/software/spm12) and our homemade MATLAB codes. The main steps were (1) removing the volumes during the first 10 seconds (14 volumes); (2) realignment of all volumes to the first volume; (3) mean-based intensity normalization (dividing each voxel by the global 4D mean value and multiplying by 1000); (4) spatial normalization to the Montreal Neurological Institute (MNI) template with EPI (2 × 2 × 2 mm voxel size) (Calhoun et al., 2017); (5) linear detrending of retaining mean; (6) bandpass filtering between 0.43 and 0.087 Hz (Termenon, Jaillard, Delon-Martin, & Achard, 2016) using the butter filter (MATLAB functions butter and filtfilt); (7) denoising: 24HMP+8Phys+Spikereg (Parkes et al., 2018), the 24HMP indicates 24 column head motion parameters (i.e., the 6 rigid-body parameter time-series, their backwards-looking temporal derivatives, plus all 12 resulting regressors squared); The 8Phys indicates 2 averaged WM and CSF signals, their temporal derivatives, plus all 4 resulting regressors squared; The Spikereg indicates spike regression, and volumes were marked as contaminated if FD_Jenk_ > 0.25 mm (Parkes et al., 2018). Here, we did not perform the slice-time correction because the TR for HCP rfMRI data is relatively short at 0.72 sec. We did not perform spatial smoothing due to its complex effects on the structure and properties of human brain networks (Alakorkko, Saarimaki, Glerean, Saramaki, & Korhonen, 2017).

#### Construct the static functional network

The first step in constructing the brain network is to define the nodes and edges. Here, for the static functional network, we utilized the Gordon parcellation with 333 regions as nodes (Gordon et al., 2016), and the Pearson correlation coefficients between pairs of nodal median time series were calculated as edges. We obtained an asymmetric connectivity matrix for each participant.

#### Construct the dynamic functional network

Here, we applied a sliding window method to construct the dynamic brain network. This framework has been enthusiastically welcomed and used repeatedly by the neuroimaging community to understand brain dynamics and their relations with cognitive abilities and different brain disorders (Preti, Bolton, & Van De Ville, 2017). To simplify, we first selected a time window with length W (from time t = 1 to time t = W), calculating the connectivity as Pearson’s correlation coefficient between each pair of time series within the time window. The window was then moved by step T, and the same calculation was repeated on the time interval [1+T, W+T]. This process was repeated until the window spans the end portion of the period. When all windows were considered, a set of connection matrices (dynamic functional networks) were obtained. Here, we used the time window with a length of 200 TRs and stepped with 17 TRs, resulting in 59 dynamic functional networks.

### Correspondence of microstructural covariance and mean functional network

#### Simple multilinear model

We used the multiple regression approach (Vazquez-Rodriguez et al., 2019) to predict the functional connections profiles of nodes based on geometric and microstructural predictors of the same node (**Fig. 1b**). The predictors were 1) path length between nodes, 2) Euclidean distance between node centroids, and 3) communicability between nodes. Binarized microstructural connectome data was used to estimate path length and communicability. Path length referred to the shortest contiguous sequence of edges between two nodes (*distance_bin.m*, BCT toolbox). Communicability (*C_ij_*) between two nodes *i* and *j* is defined as the weighted sum of all paths and walks between those nodes (*expm.m*, MATLAB 2021a built-in function). Communicability was defined for a binary adjacency matrix, A, as follows:

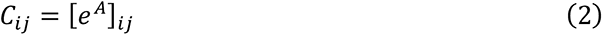

Then, we obtained the structural path length (*P_i_*), Euclidean distance (*E_i_*), and microstructural communicability (*C_i_*) for each node. We applied the regression model,

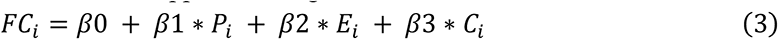

where *FC*_i_ measures the functional connections between node *i* and the other nodes. The regression coefficients *β*1, *β*2, and *β*3, as well as the intercept *β*0, were then solved by ordinary least squares (*fitlm.m*, MATLAB 2021a built-in function). The adjusted R-square for each region was termed the nodal correspondence of microstructure-function.

### Basic topology of microstructural and functional brain network

To simplify our analysis, we converted the weight networks to binary networks using a sparsity threshold value of 10% (i.e., keep the top 10% elements in the network), which has reported in previous studies (Grydeland et al., 2019; Lariviere et al., 2020). Specifically, we normalized the value in the weighted network to a range from 0 to 1. Then we extracted the minimum spanning tree (Alexander-Bloch et al., 2010) to ensure that the thresholded network is not fragmented, and the remaining values in the network were sorted by descending order and reserved in sequence until reaching the sparsity threshold value of 10%. Subsequently, we calculated four nodal metrics [clustering coefficient, local efficiency, degree centrality, global efficiency] for each network. These nodal metrics were implied with BCT toolbox (Rubinov & Sporns, 2010) (https://sites.google.com/site/bctnet). Then, we performed the hub detection, calculated the mean nodal global efficiency values across 198 participants, and defined the node with top 10% mean global efficiency values across all participants as hubs (Del Ferraro et al., 2018). For modular detection, the Louvain community detection algorithm (Newman, 2006) was applied to the mean microstructural and functional brain network for stable modular results. We determined the modularity metric (Q) at the resolution parameter, γ = 1. Q quantifies modularity, that is, Q = 0 has no more intramodular connections than expected by chance, while Q > 0 indicates a network with some community structure. We computed 10,000 iterations (function: *community_louvain.m*, BCT toolbox) and each pair of regions received an affinity score between 0 and 1. The affinity score is the times of two regions being assigned to the same module divided by the 10,000 iterations, thereby assigning higher weights to partitions with a higher modularity score. We set the affinity score ≥ 0.5 to 1 and others to 0, resulting in a binary affinity matrix, then, we then performed the modular detection on the binary affinity matrix (function: *modularity_und.m*, BCT toolbox) to obtain stable modular organization.

### Higher-order framework

To evaluate the higher-order relationships between the microstructural and functional brain networks, first, we utilized 11 non-redundant orbits and counted the orbit frequency for each node in the microstructural and functional networks with sparsity threshold 10%, separately, resulting in a *N*_nodes_ × *N*_orbits_ matrix. Second, for each node, let *f*(o*_i_*) be the frequency of orbit o*_i_* where *i* ∈ (1 … 11). Third, we calculated the normalized frequency of each *f*(o*_i_*) by dividing it with 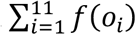, to avoid the zero in the denominator, where we added 1 to each of the frequencies, and take the logarithm (10-base) of the normalized frequency, thus the normalized orbit frequency matrix became,

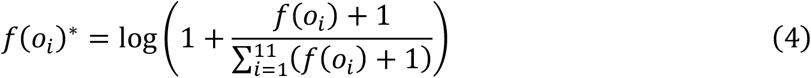

last, for the normalized orbit frequency matrix (333 × 11), we calculated the pairwise mutual information (MI) among all pairs of orbits and resulted in an 11-by-11 (*N*_orbits_ × *N*_orbits_) MI matrix, the Pearson correlation between their MI matrices was termed as the higher-order relationships of microstructural and functional brain network (**Fig. 1f-g**).

### Statistical analysis

#### Estimation of nodal microstructure variability

Here, we used the myelin content to descript the anatomical variability; for each brain region, we could obtain a 1 × 198 vector across all participants, denoted as *A*. We termed the standard deviation of each regional myelin content across all 198 participants as the variability of regional myelin,

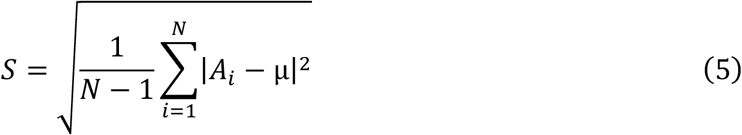

Where *N* was 198 and *μ* was the mean of *A*:

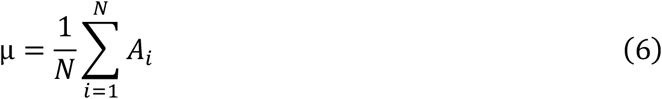

*Nodal higher-order relationships and variability*. For each participant, we counted the frequency of 11 orbits for function and myelin-based structural network and the nodal higher-order relationships were defined by the **cosine similarity** between the frequency of 11 orbits of microstructural and functional network; thus, we can obtain a higher-order relationships map for each participant at the nodal level. The standard deviation of the nodal higher-order relationships across the 198 participants was defined as the variability of the nodal higher-order relationships.

#### Estimation of the variability of nodal functional connectivity (FC)

In the current study, each participant (*P*) yielded a 333 × 333 functional connectivity matrix. For each brain region (*i*), the remaining regions’ correlation with the seed region is a 1 × 332 vector, which can be denoted as *F_n_*_-i_, we can yield a 332 (*F_n−i_* ) × 198 (*P*) matrix for all participants, denoted as *M_i_.* We termed the functional connectivity variability of each brain region as *V_region(i)_* = 1 − *ICC(M_i_)*. The intraclass correlation coefficient (ICC) could be used to define the stability of scores over participants,

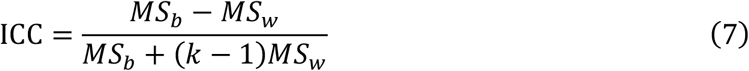

where *MS_b_* was the between-subject mean square, *MS_w_* was the within-subject mean square, *k* was the number of repeated observations for each participant (198 here).

## APPENDIX

### Correlation between microstructural and functional centrality and nodal higher-order microstructure-function correspondence

Correlation analysis (Pearson correlation) was performed to examine the relationships between the *mean higher-order microstructure-function correspondence* values with the *mean functional centrality* and *mean microstructural centrality* across all participants. In both cases the correlations were moderately and significant (all *P-values* < 0.005, *FDR correction*), suggesting that nodal higher-order microstructure-function correspondence can partially be explained by microstructural or functional nodal centrality.

**Fig. S1.**
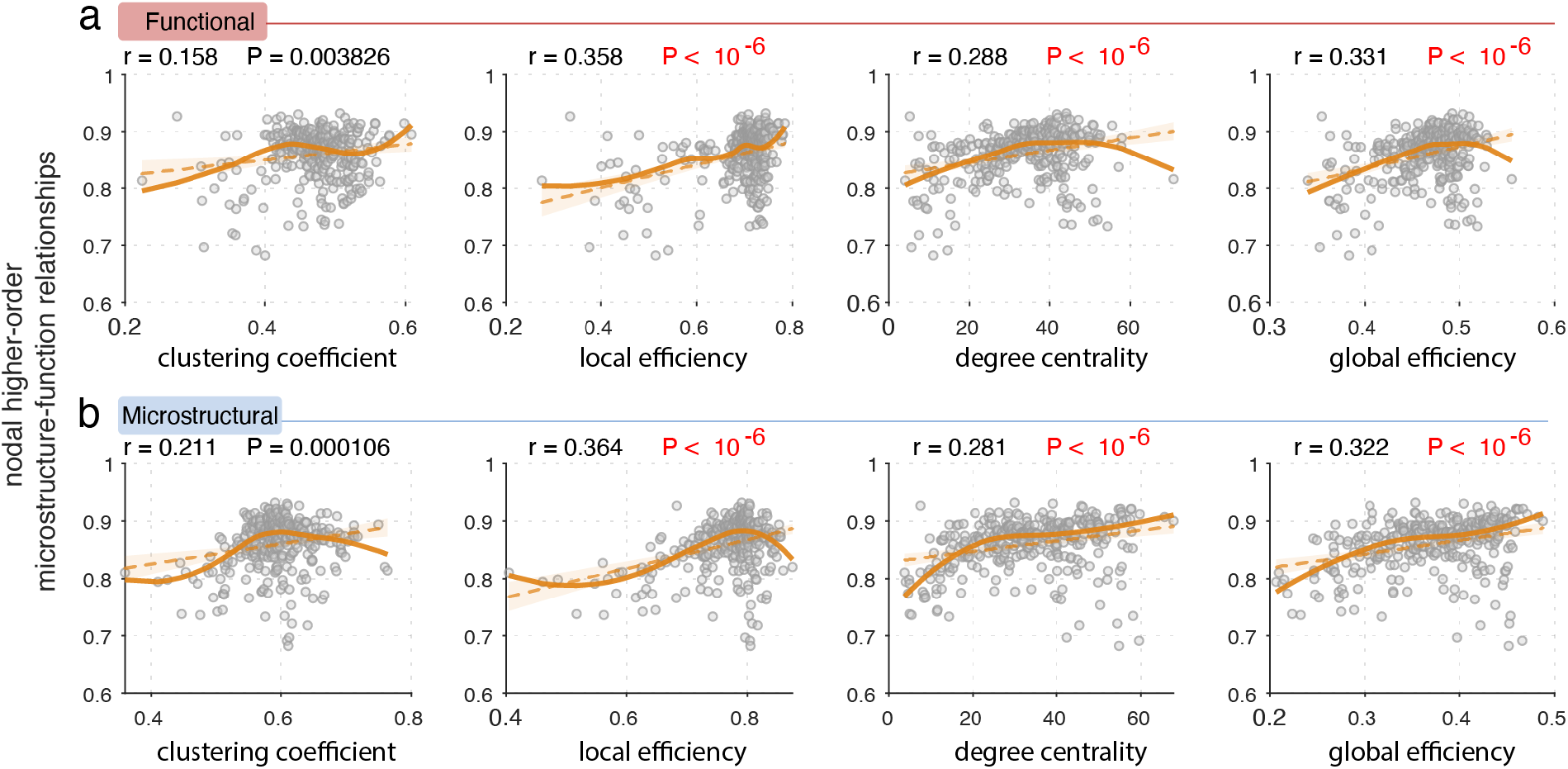
| Correlation between microstructural and functional centrality and microstructure–function correspondence. (**a**) Correlation between mean nodal higher-order microstructure-function correspondence and mean functional nodal centrality. (**b**) Correlation between mean nodal higher-order microstructure-function correspondence and mean microstructural nodal centrality. The dashed line indicates linear (least squares) fit; the solid line indicates non-parametric regression [LOESS (locally estimated scatterplot smoothing)] fit.

### Local (circuit-level) higher-order microstructure-function relationship

**Fig. S2.**
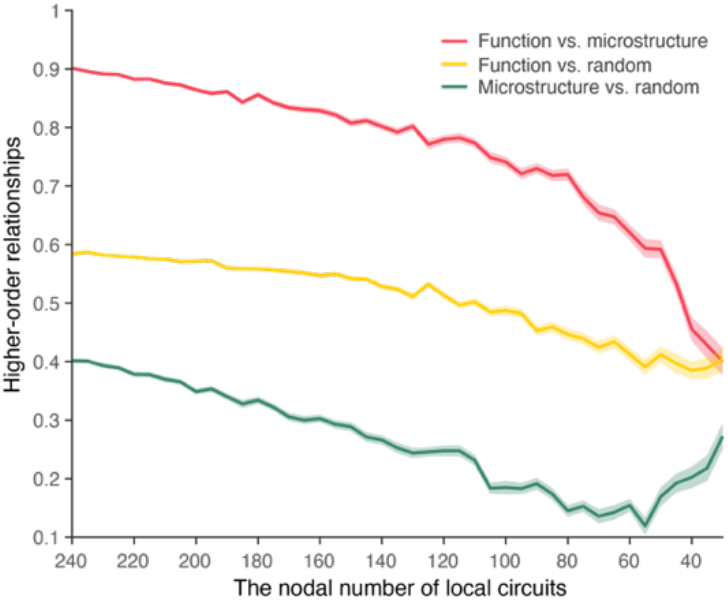
| Higher-order relationships across different subnetworks/scales. Higher-order relationships of microstructure- function are larger than their relations with random networks from 240 nodes to 30 nodes.

### Additional analysis

#### Applying the higher-order method to network classification

We predict that higher-order similarity between networks within the same category will be greater than between networks within different categories since the higher-order framework may capture the hidden information well. Three types of networks (functional, microstructural, and random), each type containing 396 networks (198 networks with sparsity 0.1 and 198 networks with sparsity 0.2), are used to prove our hypothesis. Any paired network’s higher-order interactions are calculated, producing an 1188 by 1188 similarity matrix (**Fig. S3a**). The higher-order similarity of any two categories of networks is shown in (**Fig. S3b**), the networks of the same category exhibit stronger higher-order similarity than different categories of networks. Positive class refers to similarities between networks of the same category, and negative class refers to similarities between different types of networks. According to the receiver operating characteristic (ROC) curve analysis, the higher-order property makes clustering networks of similar types more sensitive, with an AUC of 0.998 and an Area Under Precision-Recall curve (AUPR) of 0.996. In contrast, none of the lower-order measures is above 0.9 (**Fig. S3c-d**), highlighting the importance of high-order properties for clustering.

#### Relation between nodal metrices and orbits

For a better understanding of the orbits, the Spearman correlation analysis is conducted between the four nodal metrics and the normalized nodal frequency of 11 orbits, resulting in a 4 by 11 matrix for each participant. K-means clustering is used to identify potential categories. The classification of functional networks is similar to that of microstructural networks (**Fig. S3e-f**). The clustering coefficient and local efficiency are in one cluster, and the degree centrality and global efficiency are in another cluster. Hence, one cluster may reflect local properties while another owns global properties. For the 11 orbits, the O1, O2, O4, O6, and O9 is grouped together, while the O3, O5, O7, O8, O10, and O11 were grouped together (**Fig. S3e-f**). Intriguingly, O1, O2, O4, O6, and O9 correspond to the peripheral, their degree centrality is 1. In contrast, O3, O5, O7, O8, O10, and O11 are the most likely clusters or hubs, with degree centrality of at least two (**Fig. 3b** *in main text*). Using the higher-order framework, we can determine the interactions between different functional subnetworks or create network of networks to describe the relationships among complex systems for other disciplines.

**Fig. S3.**
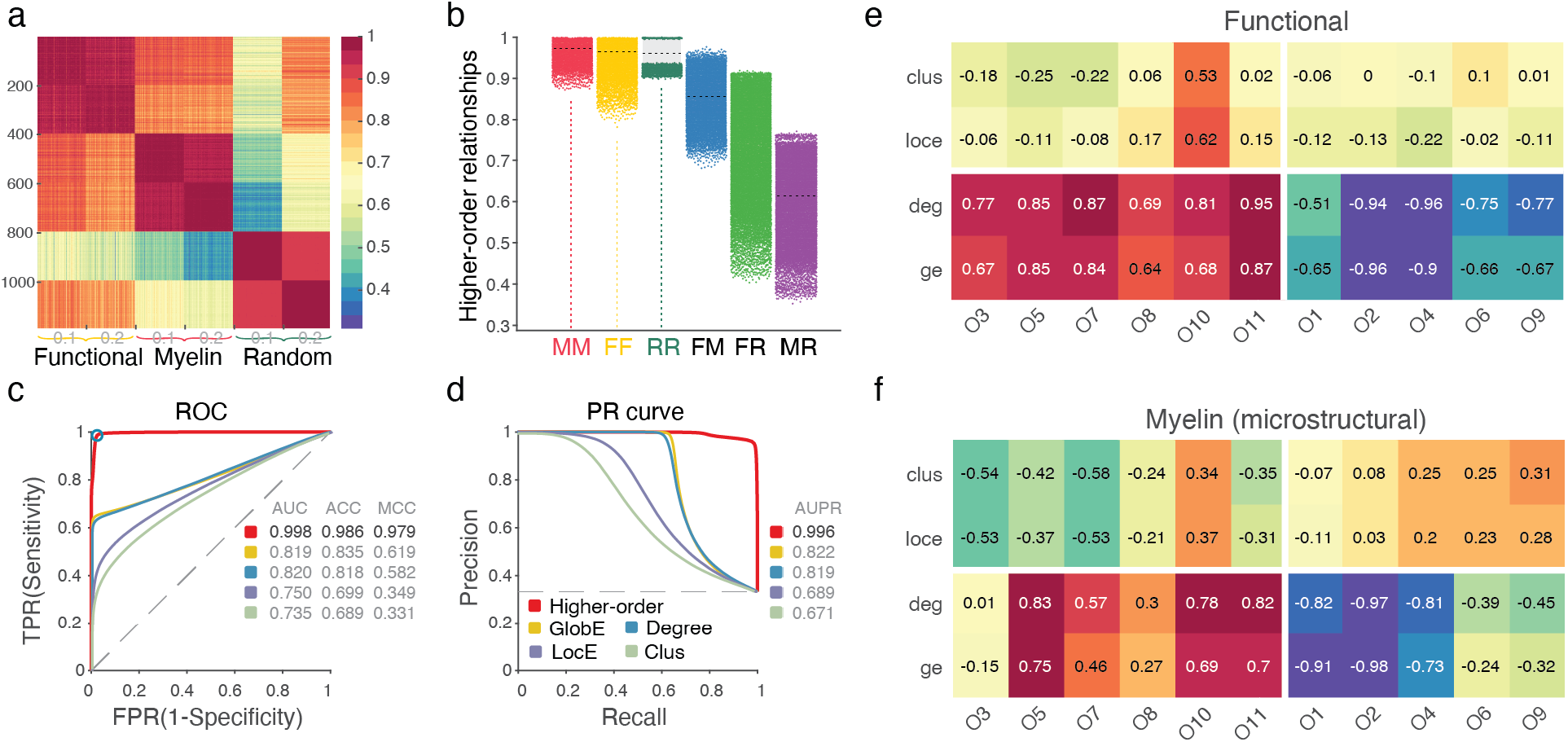
| Network classification and the relationship between nodal metrics and orbits. (**a**) The higher-order relationship matrix of 1,088 networks: 396 functional, 396 microstructural, and 396 random networks. Two sparsity values of 0.1 and 0.2 are used, under each sparsity value, there are 198 networks. (**b**) The higher-order similarity between different networks. MM indicates microstructural & microstructural; FF indicates functional & functional; RR indicates random & random; FM represents functional & microstructural; FR represents functional & random; MR represents microstructural & random. (**c**) We define the higher-order similarity values coming from the same type of networks as positive class (MM, FF, RR) and the similarity values coming from different types of networks as the negative class (FM, FR, MR), and we utilize the ROC curve analysis to measure the quality of higher-order classification for the 1188 networks. ROC curves, which plot the true positive rate (TPR) versus the false positive rate (FPR), and (**d**) PR curves, which plot the precision versus the recall. True positives (TP); false positives (FP); false negatives (FN); and true negatives (TN). The accuracy (ACC) is equal to (TP+TN) / (all samples), and the Matthews correlation coefficient (MCC) is equal to 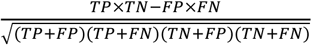. The results show that the higher-order property exhibits an excellent quality in classifying the networks compared to the lower-order property. GlobE, global efficiency; Degree, degree centrality; LocE, local efficiency; Clus, clustering coefficient. (**e-f**) The relationship between the nodal metrics and normalized orbits’ frequency. The clustering results show a similar pattern in both networks. For nodal metrics, the clustering coefficient and local efficiency are grouped together, while the degree centrality and global efficiency are classified into a group. For orbits, the O1, O2, O4, O6, and O9 were grouped together, and the O3, O5, O7, O8, O10, and O11 were grouped together.

### Robustness

The robustness and reproducibility of the main findings are vital for scientific research. Here, we also evaluate our main results against several factors, including (I) the effect of sparsity thresholding; (II) the effect of sample size; (III) split-half reliability for nodal higher-order interactions; (IV) higher-order relationships on other different types of networks; (V) the effect of different embedding methods; (VI) the reliability of constructing the individual microstructural network.

#### Effect of sparsity thresholding

Despite advances in the human brain connectome, the selection of sparsity values is still under debate. First, we also evaluated whether our main results could be replicated using an efficiency cost optimization (ECO) filter threshold strategy (De Vico Fallani, Latora, & Chavez, 2017), which has approximately 3/(*N*-1). *N* indicates the number of nodes, here *N* = 333. Thus, the sparsity value was 9 ‰. The result suggests a similar pattern compared with the sparsity value of 10% (**Fig. S4a**). We also use the mean microstructural and functional network to test the changes of higher-order interactions across the sparsity from 0.02 to 0.4 with an interval of 0.02 (**Fig. S4b**). We find that the higher-order microstructure-function interactions can distinguish from the random networks with a lower sparsity value, less than 0.13 (**Fig. S4c**). Furthermore, we perform the correlation analysis between higher-order interactions with individual personality scores and detect the difference of higher-order interactions between three age groups (**Fig. S5**).

**Fig. S4.**
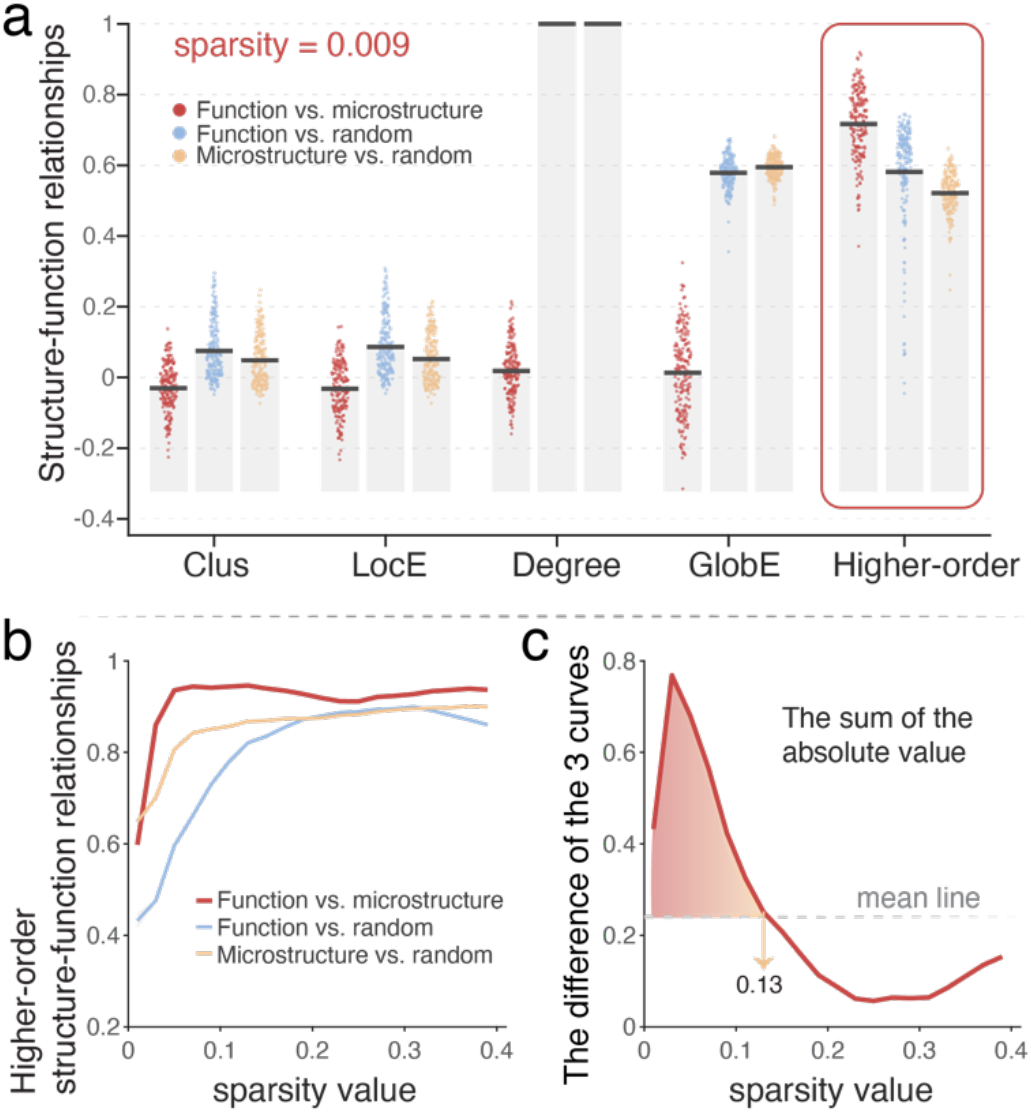
| Effect of sparsity thresholding. (**a**) Lower- and higher-order structure-function relationships under the sparsity value of 9 ‰. The microstructural and functional networks show a minimum degree of correlation at the lower-order level and exhibit a higher similarity at the higher-order level. At the lower-order level, the microstructural and functional networks exhibit a lower correlation compared with their corresponding random networks. At the higher-order level, the microstructural and functional networks exhibit enhanced correlation than their relationships with corresponding random networks. Clus, clustering coefficient; LocE, local efficiency; Degree, degree centrality; GlobE, global efficiency; Higher-order, higher-order relationships. (**b**) Higher-order interactions across the sparsity value from 0.02 to 0.4, we can observe that the higher-order microstructure-function interactions are stronger than their corresponding random networks. (**c**) The differences among the three curves are shown in **Fig. S4b,** results show that the higher-order interaction owns a solid ability to differentiate under the sparsity value of 0.13.

**Fig. S5.**
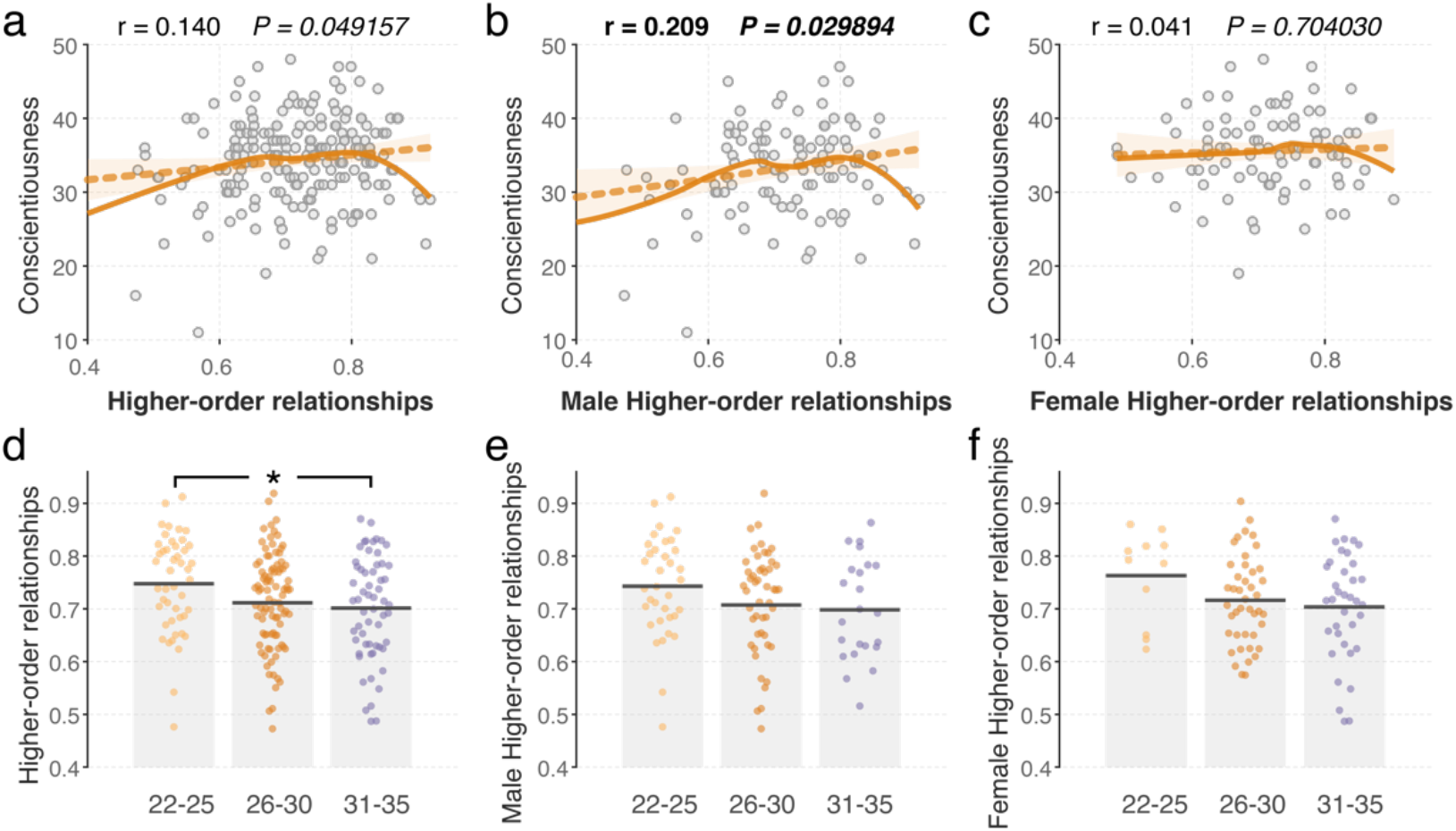
| Higher-order relationships and association with cognition. (**a**) At the sparsity value of **9 ‰**, the higher-order microstructure-function interactions are positively correlated with conscientiousness, (**b**) especially for male participants, (**c**) no significant results were observed for female group. (**d**) The higher-order relationships show a decreased pattern from the 22-25 group to the 31-35 group. No significant results were observed for male (**e**) or female participants (**f**).

#### Effect of sample size

To verify whether our main results are stable with a changeable sample size, we randomly select 100 participants from 198 participants at one time (10,000 times sampling), and then recalculate the relationship between higher-order similarity and personality scores, and the change with different age groups. The result for one-time sampling is shown in **Fig. S6a-b**; we can observe the higher-order interactions are significantly negatively correlated with agreeableness score for male participants, and show a declining trend across the age groups. The results of 10,000 times sampling, showing the higher-order interactions is decline trend across the age groups for especially for the male participants (**Fig. S6c**), and the correlations between higher-order interactions and agreeableness score for male participants are at least a 50% chance of being significant (median r = -0.269, median *P*-value = 0.048), see **Fig. S6d**.

When we randomly select 150 participants from 198 participants at one time (10,000 times sampling), the correlations between higher-order interactions and agreeableness score for male participants are 96% chance (the number of times *P* greater than 0.05 is 400, the probability is 400/10,000 = 0.04) of being significant (median r = -0.268, median *P* value = 0.012). This shows that the current sample size is enough to detect a significant correlation between higher-order relationship and personality scores.

#### Split-half reliability for nodal higher-order interactions

To examine the robustness of nodal higher-order interactions, we split all 198 subjects into two subgroups; one is the odd group (1, 3, 5, …,197), the other one is the even group (2, 4, 6, …, 198). Then, we perform the Pearson correlation in the two subgroups; a high correlation coefficient between the two subgroups is observed (r = 0.932, *P* < 10^-6^), indicating high reliability (**Fig. S6e)**.

**Fig. S6.**
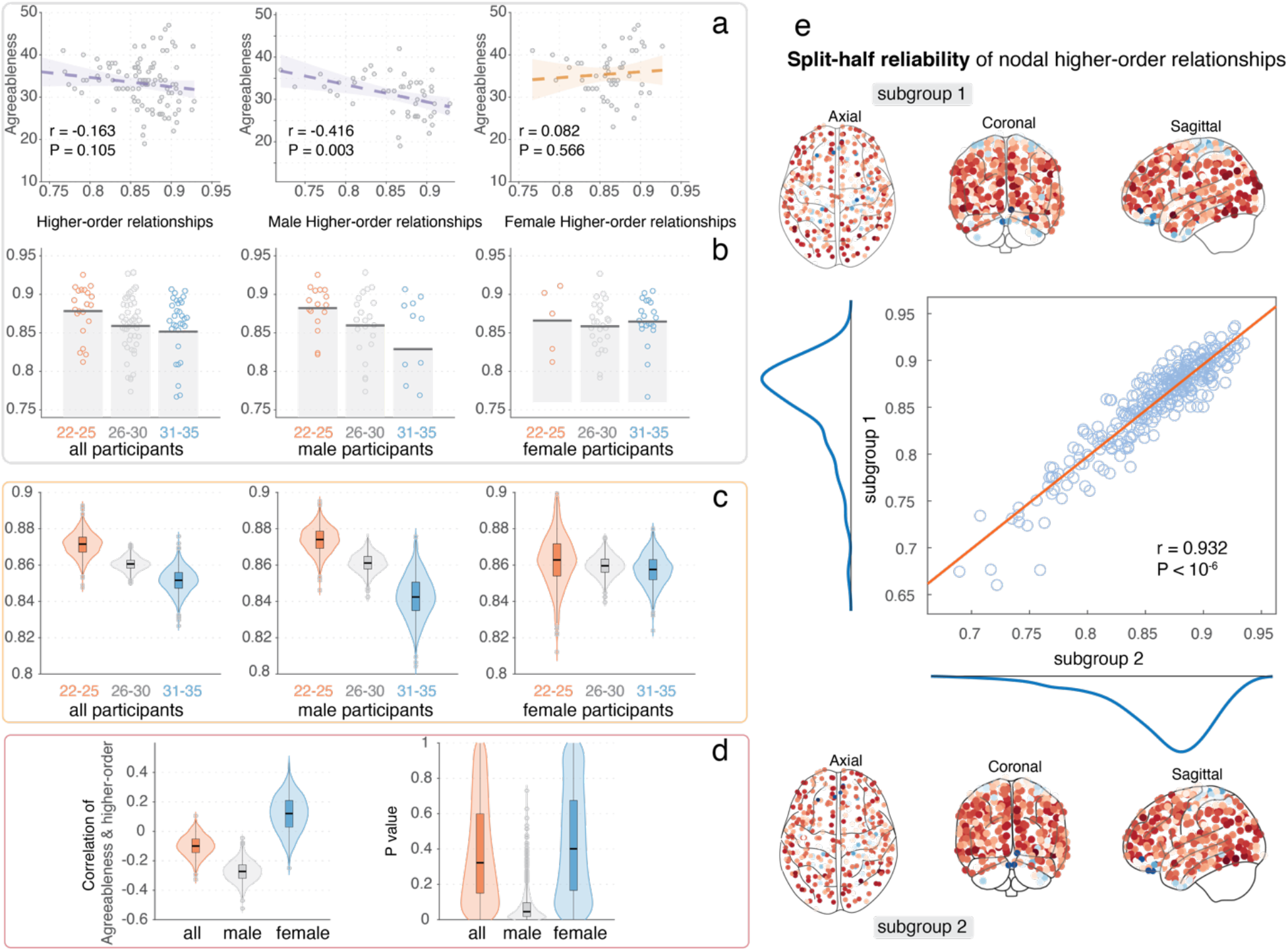
| Effect of sample size and split-half reliability. (**a-b**) At the sparsity value of 0.1, we randomly selected 100 participants from all 198 participants, and the results for one time sample are shown. (**c-d**) The mean results of 10,000 samplings for higher-order microstructure-function relationship with cognition scores and gender. (e) Split-half reliability for nodal higher-order interactions, subgroup 1 and subplot 2 have high correlation.

#### Higher-order relationships in different types of network

To further evaluate the performance of the higher-order framework. We utilize two widely used network models: small-world network and Barabási-Albert network. First, we evaluate the changes in higher-order relationships from regular network to random networks. We apply the Watts-Strogatz model to construct a serial network from regular to random networks. There are two basic stages: 1) Create a circle lattice with 2000 nodes of average degree 100. 2) For each edge in the graph, rewiring the target node with probability *P* (0 to 1, the interval is 0.01). The rewired edge cannot be duplicated or self-loop. When *P* = 0, the model yields a ring lattice. Alternatively, the ring lattice is converted into a random graph when *P* = 1. Thus, we obtain 101 networks and computed the higher-order relationships for any paired networks, resulting in a 101 × 101 matrix. We observe that the same type of networks exhibited strong higher-order relationships (e.g., regular vs. regular or random vs. random) and different types of networks (e.g., regular vs. random) exhibited lower higher-order relationships, see **Fig. S7a**. Second, we want to check the higher-order relationships among different types of networks with different network sizes (the number of nodes in the network).

For the small-world network model and the Barabási-Albert network model, we simulate 20 networks with network size from 100 to 2000 (the interval is 100) and the average degree is 20, for each network model. Then we calculate the higher-order relationships for any paired network. Similar results are observed, higher-order relationships in the same type of network, even with different network size, exhibit higher relationships than different type networks; see **Fig. S7b**. Third, we also want to evaluate the performance of higher-order relationships on real brain networks with different network sizes. Here, we randomly select a participant and used the *Schaefer Parcellations 7 network* ranging from 100 to 1000 parcels (100, 200, 300, 400, 500, 600, 800, 1000) to construct individual microstructural and functional brain networks. Then we calculate the higher-order relationships between any pair of the 16 networks. A similar pattern is observed with the main findings; see **Fig. S7c** for details.

**Fig. S7.**
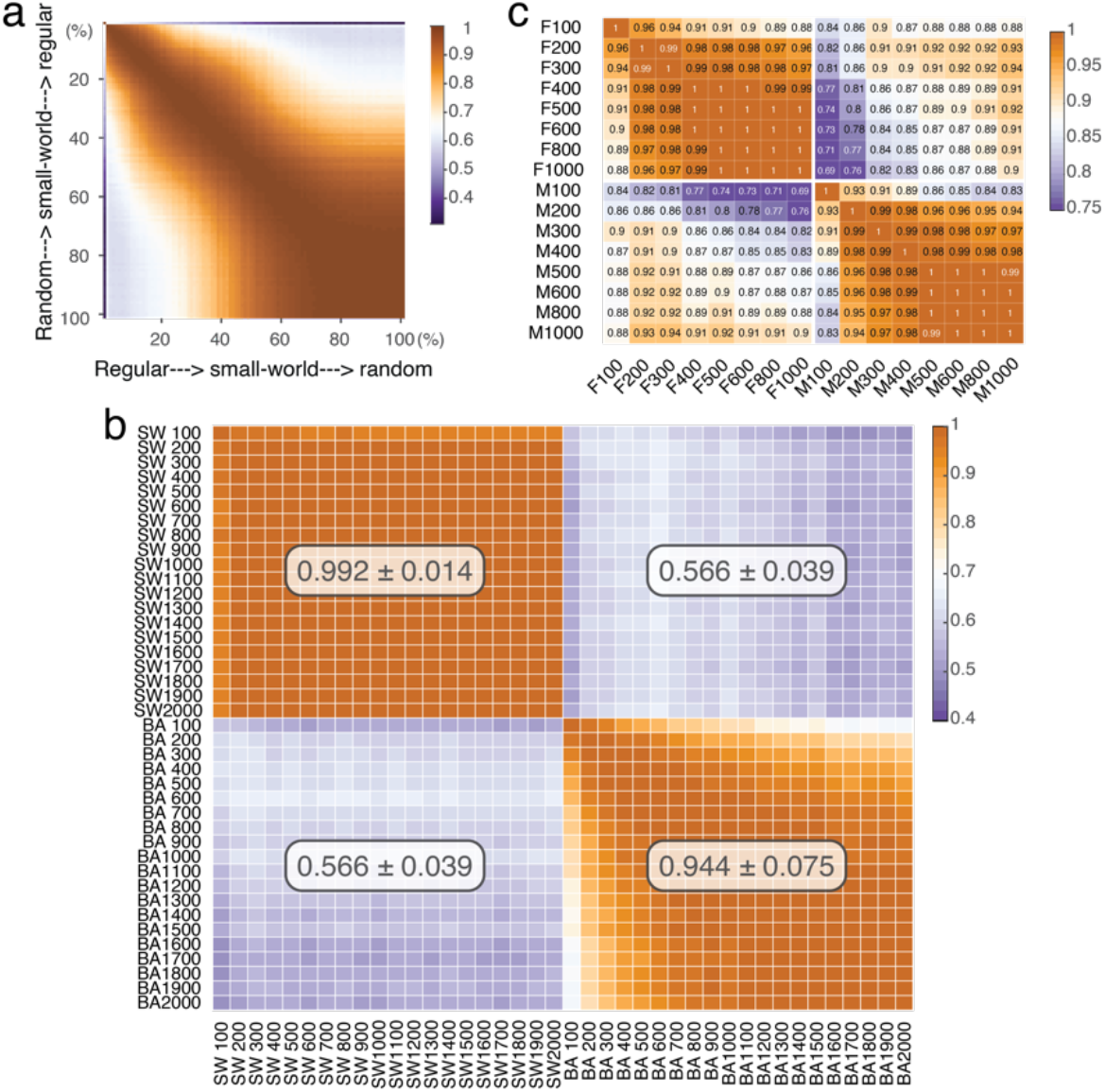
Higher-order relationships across different networks. (**a**) Higher-order relationships across regular, small-world, and random networks. (**b**) Higher-order relationships between small-world (SW) and Barabási–Albert (BA) networks with different network sizes. SW 100 indicates SW network with 100 nodes; BA 100 indicates BA network with 100 nodes, and so on. (**c**) Higher-order relationships between microstructural and functional networks with different network sizes, the images of ***subject ID: 102109*** were used. F: functional network; M: microstructural network; F100 indicates functional network with 100 nodes; M100 indicates microstructural network with 100 nodes, and so on.

#### Effect of different embedding methods

In addition to using the orbit embedding, we also evaluate the effect of the nodal neighborhood embedding framework. Specifically, first, we thresholded the network with a 10% sparsity value, i.e., to keep the top 10% edges, then, we identified the **shortest path matrix** of the thresholded network, here, we consider the 1st to 6th step neighbors for each node, and we calculate the number of neighbors for each node from the 1st step to 6th step, obtain a N × 6 matrix, N is the number of nodes. Then we calculate the

Euclidean distance for any paired step neighbors within the N × 6 matrix, resulting in a 6 × 6 Euclidean distance matrix. Thus, for any two networks, we could compress each of them into a 6 × 6 Euclidean distance matrix, the higher-order relationship was termed as the Pearson correlation of two Euclidean distance matrices. The workflow of this neighbor embedding method can be seen in **Fig. S8.** Here, we explore the higher-order relationships by 6 step nodal neighborhood embedding for functional, microstructural, and random networks. We observe significantly higher relationships between microstructural and functional networks than their relationships with random networks (**Fig. S9a**) and exhibited a decreased trend across three age groups (**Fig. S9b**). However, we do not observe any significant correlation between the higher-order relationships and individual personality (Big-Five) scores using the *neighborhood embedding framework*.

**Fig. S8.**
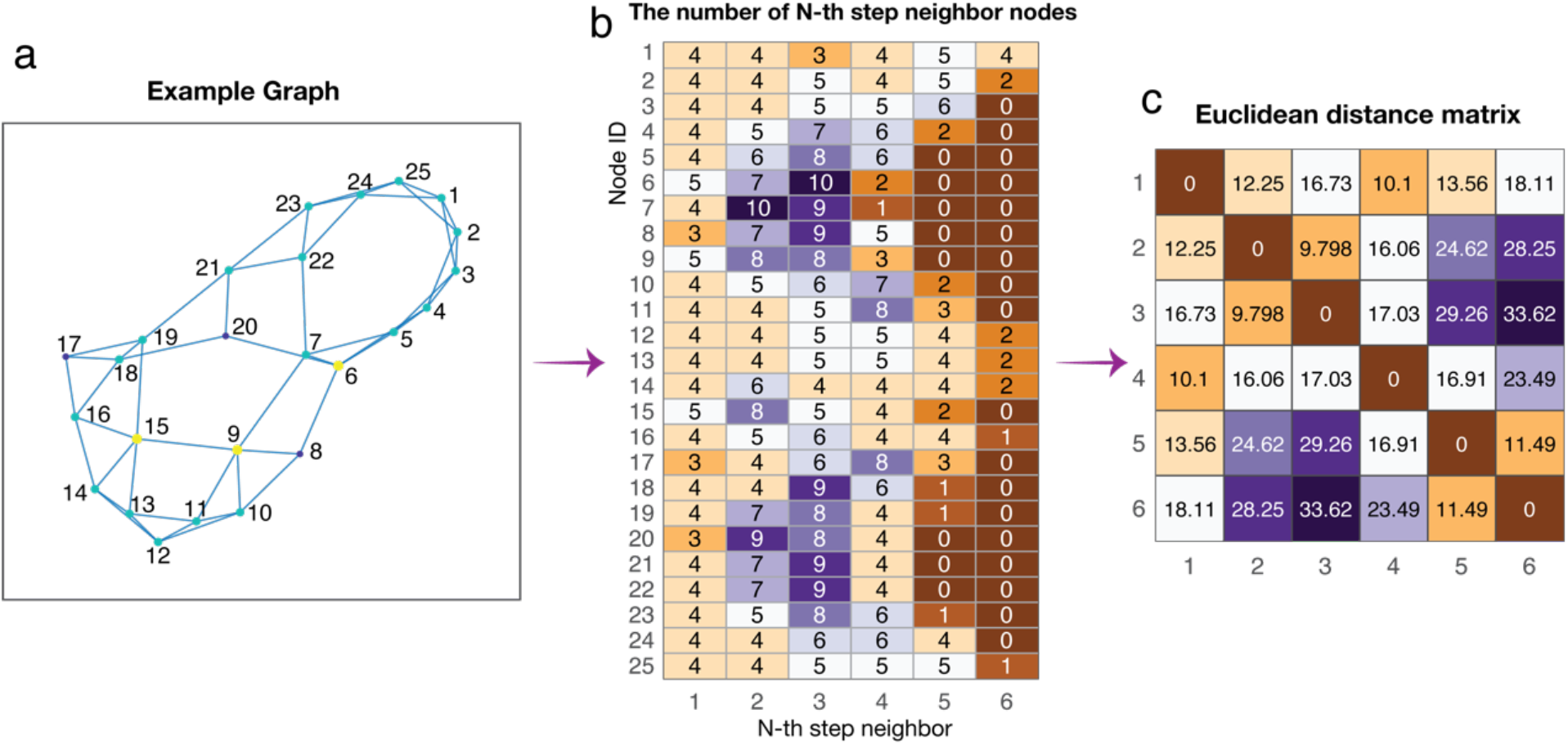
Workflow of the neighborhood embedding method. (**a**) Example graph with 25 nodes and 50 edges. (**b**) Up to six-step nodal neighborhood embedding, counting the number of neighbors for each step for each node, the shortest path distance was used to depict the number of neighbors in each step. (**c**) Euclidean distance matrix of the 6 features embeddings.

**Fig. S9.**
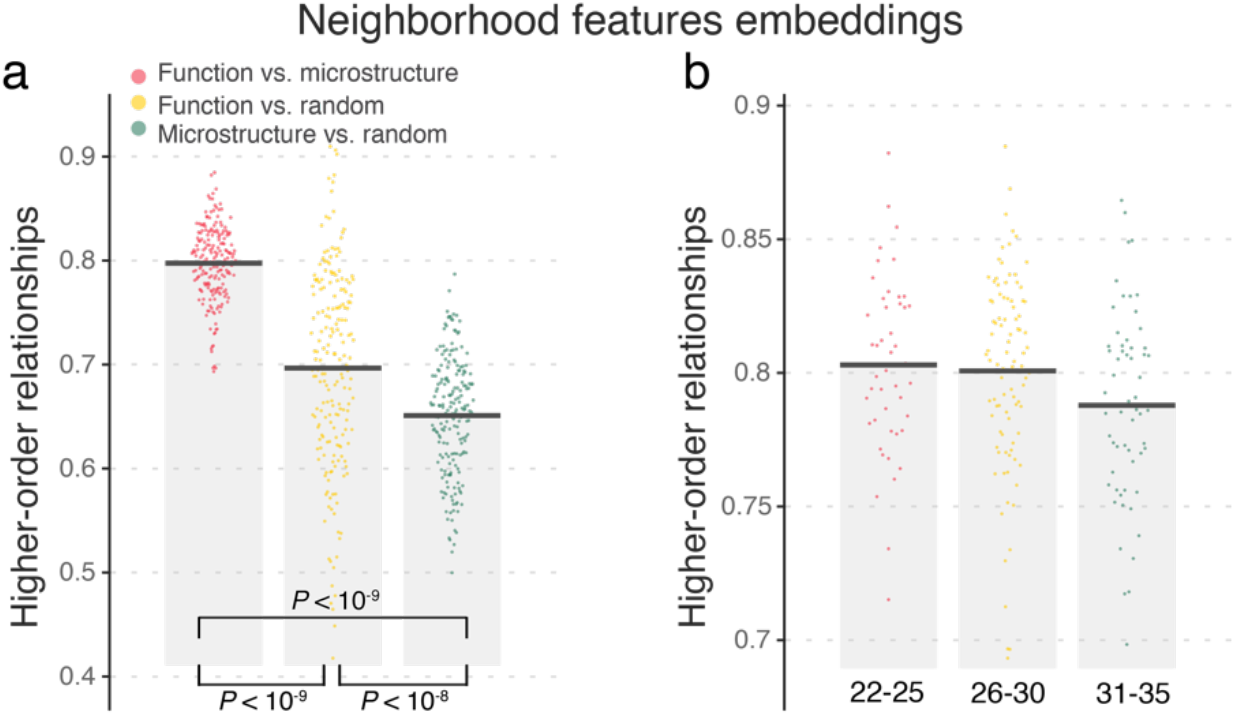
Higher-order relationships by neighborhood embedding method. (a) Higher-order microstructure-function relationships are higher than their relations with random networks. (b) Microstructure-function higher-order relationships show decreased trend across three age groups (no significant group differences were observed).

#### Reliability of constructing the individual microstructural network

In addition, we also evaluate the reliability of our method for constructing the individual microstructural network, another HCP test-retest dataset that included 20 participants is used to perform the intraclass correlation coefficient (ICC) analysis. The ID information of 20 participants can be seen in **Table S2**. The connection matrix exhibited high ICC (0.960±0.016), we thresholded these networks with a sparsity of 0.1 and calculate the nodal degree centrality and global efficiency for each network. For nodal metrics, nodal degree exhibits a high ICC (0.911±0.079), nodal global efficiency exhibits a high ICC (0.936±0.059), and these regions with high ICC values are mainly located in the parietal and occipital lobes, while the regions within the default mode network show a lower ICC. See **Fig. S10**.

**Fig. S10.**
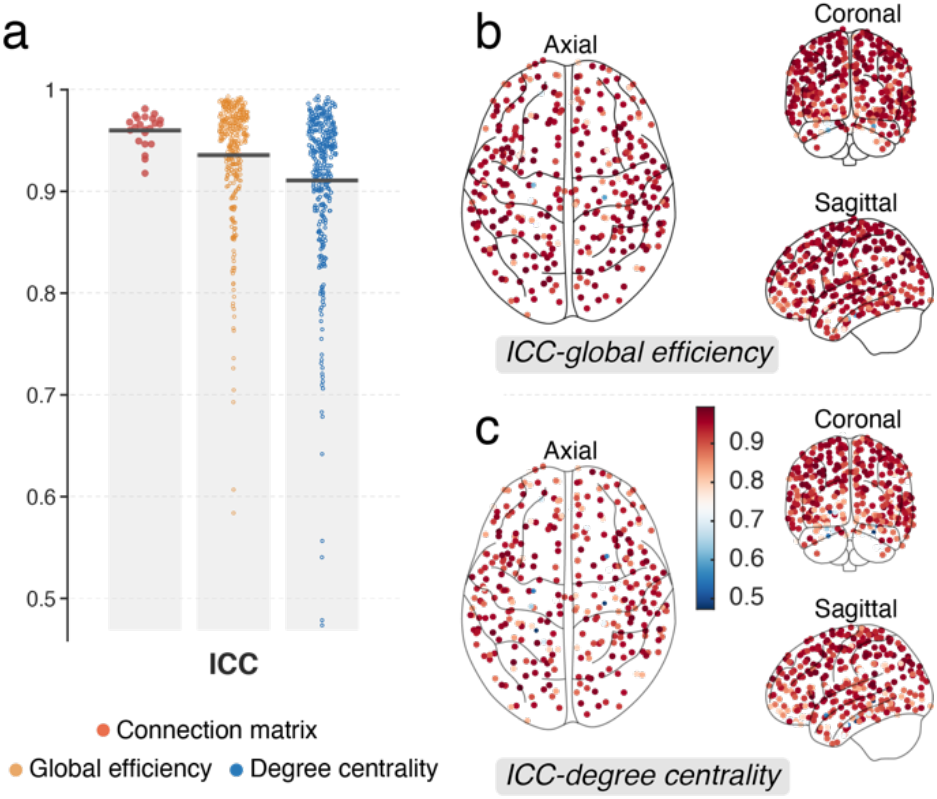
The reliability of our method for constructing the individual microstructural network. (**a**) The results indicate that the connection matrix, degree centrality, and global efficiency exhibit high ICC values. (**b**) The ICC map of global efficiency. (**c**) The ICC map of degree centrality. We observe the occipital and parietal lobes have a higher ICC value.

Collectively, the current findings are robust to some factors and the method for constructing individual myelin-based microstructural network is highly reliable.

**Table S1.** We download unprocessed MR data of **213** participants from the “S1200” release of the Human Connectome Project (HCP). Eight participants who have a larger **head movement** were excluded (IDs: 161832, 192237, 362034, 392447, 453542, 468050, 804646, 962058). Seven participants ≥ **36 years old** were further discarded (IDs: 171128, 202820, 209531, 689470, 757764, 902242, 905147).

|  | SUBJECT | RELEASE | ACQUISITION | GENDER | AGE | NEO_A | NEO_O | NEO_C | NEO_N | NEO_E |
| --- | --- | --- | --- | --- | --- | --- | --- | --- | --- | --- |
| 1 | 102109 | S1200 | Q13 | M | 26-30 | 39 | 41 | 30 | 19 | 37 |
| 2 | 102614 | S1200 | Q12 | M | 22-25 | 34 | 30 | 42 | 5 | 35 |
| 3 | 102715 | S1200 | Q12 | M | 26-30 | 38 | 26 | 38 | 6 | 36 |
| 4 | 103010 | S1200 | Q12 | M | 22-25 | 29 | 33 | 40 | 19 | 28 |
| 5 | 103212 | S1200 | Q13 | M | 31-35 | 36 | 31 | 21 | 17 | 32 |
| 6 | 106824 | S1200 | Q13 | M | 22-25 | 28 | 29 | 34 | 22 | 29 |
| 7 | 108020 | S1200 | Q12 | M | 22-25 | 29 | 17 | 33 | 16 | 34 |
| 8 | 111211 | S1200 | Q12 | F | 26-30 | 33 | 27 | 32 | 20 | 36 |
| 9 | 113316 | S1200 | Q12 | M | 22-25 | 30 | 31 | 37 | 12 | 29 |
| 10 | 113417 | S1200 | Q12 | F | 26-30 | 32 | 25 | 39 | 19 | 32 |
| 11 | 114116 | S1200 | Q12 | F | 26-30 | 32 | 23 | 30 | 28 | 33 |
| 12 | 115724 | S1200 | Q12 | F | 22-25 | 33 | 21 | 36 | 19 | 32 |
| 13 | 116423 | S1200 | Q09 | M | 26-30 | 26 | 33 | 29 | 20 | 35 |
| 14 | 117021 | S1200 | Q12 | F | 26-30 | 29 | 34 | 31 | 20 | 18 |
| 15 | 117728 | S1200 | Q10 | F | 22-25 | NaN | NaN | NaN | NaN | NaN |
| 16 | 118831 | S1200 | Q13 | M | 22-25 | 25 | 33 | 31 | 39 | 23 |
| 17 | 119025 | S1200 | Q12 | M | 26-30 | 29 | 27 | 33 | 1 | 11 |
| 18 | 120010 | S1200 | Q13 | F | 26-30 | 35 | 39 | 42 | 8 | 30 |
| 19 | 120414 | S1200 | Q12 | F | 26-30 | 34 | 32 | 34 | 12 | 30 |
| 20 | 122418 | S1200 | Q13 | F | 26-30 | 34 | 23 | 36 | 15 | 36 |
| 21 | 123723 | S1200 | Q09 | F | 31-35 | 25 | 22 | 31 | 34 | 34 |
| 22 | 125222 | S1200 | Q12 | M | 26-30 | 28 | 35 | 33 | 24 | 32 |
| 23 | 125424 | S1200 | Q11 | F | 31-35 | 38 | 10 | 47 | 11 | 27 |
| 24 | 126426 | S1200 | Q12 | F | 31-35 | 38 | 20 | 38 | 22 | 26 |
| 25 | 127226 | S1200 | Q08 | M | 22-25 | 33 | 30 | 36 | 16 | 30 |
| 26 | 127731 | S1200 | Q09 | F | 31-35 | 34 | 29 | 36 | 15 | 21 |
| 27 | 127832 | S1200 | Q09 | F | 31-35 | 46 | 24 | 37 | 2 | 47 |
| 28 | 130114 | S1200 | Q08 | M | 26-30 | 32 | 44 | 37 | 3 | 40 |
| 29 | 130518 | S1200 | Q07 | F | 31-35 | 33 | 28 | 40 | 14 | 29 |
| 30 | 130720 | S1200 | Q13 | M | 31-35 | 25 | 32 | 31 | 21 | 20 |
| 31 | 134627 | S1200 | Q09 | M | 26-30 | 39 | 29 | 29 | 8 | 29 |
| 32 | 135124 | S1200 | Q13 | F | 31-35 | 38 | 31 | 25 | 26 | 37 |
| 33 | 135629 | S1200 | Q12 | M | 22-25 | 29 | 26 | 29 | 26 | 32 |
| 34 | 136126 | S1200 | Q09 | M | 26-30 | 42 | 23 | 32 | 11 | 28 |
| 35 | 136631 | S1200 | Q09 | M | 22-25 | 32 | 18 | 31 | 17 | 34 |
| 36 | 137431 | S1200 | Q13 | F | 31-35 | 43 | 26 | 47 | 1 | 33 |
| 37 | 137532 | S1200 | Q12 | M | 31-35 | 29 | 25 | 32 | 15 | 35 |
| 38 | 138130 | S1200 | Q09 | F | 31-35 | 30 | 35 | 32 | 24 | 29 |
| 39 | 138332 | S1200 | Q13 | M | 26-30 | 34 | 32 | 38 | 10 | 45 |
| 40 | 139435 | S1200 | Q13 | F | 26-30 | 34 | 18 | 37 | 11 | 26 |
| 41 | 143224 | S1200 | Q12 | F | 26-30 | 35 | 25 | 36 | 18 | 30 |
| 42 | 143830 | S1200 | Q06 | F | 26-30 | 27 | 22 | 29 | 23 | 31 |
| 43 | 144933 | S1200 | Q12 | M | 26-30 | 27 | 34 | 26 | 38 | 32 |
| 44 | 145632 | S1200 | Q08 | M | 26-30 | 24 | 29 | 16 | 41 | 28 |
| 45 | 146735 | S1200 | Q12 | F | 31-35 | 31 | 25 | 40 | 16 | 24 |
| 46 | 146836 | S1200 | Q11 | M | 26-30 | 32 | 29 | 31 | 19 | 31 |
| 47 | 147636 | S1200 | Q10 | F | 22-25 | 37 | 29 | 27 | 21 | 18 |
| 48 | 151021 | S1200 | Q09 | M | 31-35 | 36 | 21 | 38 | 11 | 30 |
| 49 | 151324 | S1200 | Q09 | M | 22-25 | 21 | 35 | 31 | 27 | 31 |
| 50 | 151930 | S1200 | Q12 | M | 31-35 | 33 | 29 | 45 | 20 | 26 |
| 51 | 152225 | S1200 | Q10 | M | 31-35 | 32 | 23 | 38 | 1 | 25 |
| 52 | 152427 | S1200 | Q13 | F | 31-35 | 35 | 28 | 31 | 27 | 20 |
| 53 | 153126 | S1200 | Q13 | F | 26-30 | 29 | 25 | 34 | 16 | 27 |
| 54 | 153934 | S1200 | Q12 | M | 26-30 | 36 | 22 | 34 | 4 | 35 |
| 55 | 154330 | S1200 | Q04 | M | 26-30 | 27 | 29 | 31 | 10 | 34 |
| 56 | 165436 | S1200 | Q12 | M | 26-30 | 37 | 30 | 22 | 28 | 28 |
| 57 | 165941 | S1200 | Q09 | F | 26-30 | 23 | 23 | 29 | 22 | 22 |
| 58 | 167440 | S1200 | Q13 | F | 26-30 | 44 | 35 | 38 | 20 | 25 |
| 59 | 168947 | S1200 | Q12 | F | 26-30 | 35 | 35 | 34 | 13 | 35 |
| 60 | 169545 | S1200 | Q13 | M | 22-25 | 28 | 26 | 35 | 12 | 36 |
| 61 | 171734 | S1200 | Q13 | F | 26-30 | 39 | 20 | 36 | 11 | 33 |
| 62 | 172635 | S1200 | Q12 | M | 31-35 | 25 | 35 | 30 | 26 | 33 |
| 63 | 175136 | S1200 | Q12 | F | 22-25 | 31 | 24 | 39 | 12 | 34 |
| 64 | 176845 | S1200 | Q11 | F | 26-30 | 47 | 28 | 29 | 8 | 36 |
| 65 | 177140 | S1200 | Q13 | F | 31-35 | 38 | 32 | 35 | 16 | 26 |
| 66 | 180230 | S1200 | Q13 | F | 26-30 | 37 | 36 | 36 | 7 | 36 |
| 67 | 180533 | S1200 | Q12 | F | 31-35 | 37 | 27 | 36 | 15 | 32 |
| 68 | 183741 | S1200 | Q12 | M | 31-35 | 37 | 37 | 34 | 23 | 24 |
| 69 | 185038 | S1200 | Q11 | M | 31-35 | 37 | 31 | 33 | 25 | 25 |
| 70 | 186040 | S1200 | Q09 | M | 26-30 | 37 | 35 | 32 | 14 | 34 |
| 71 | 186545 | S1200 | Q13 | F | 31-35 | 32 | 23 | 32 | 23 | 28 |
| 72 | 186848 | S1200 | Q12 | F | 26-30 | 38 | 33 | 36 | 15 | 40 |
| 73 | 186949 | S1200 | Q13 | M | 22-25 | 32 | 18 | 45 | 14 | 40 |
| 74 | 188145 | S1200 | Q12 | F | 31-35 | 39 | 21 | 39 | 12 | 24 |
| 75 | 189652 | S1200 | Q13 | M | 26-30 | 43 | 13 | 35 | 29 | 36 |
| 76 | 191235 | S1200 | Q10 | M | 22-25 | 34 | 20 | 38 | 2 | 30 |
| 77 | 193845 | S1200 | Q13 | M | 22-25 | 37 | 28 | 42 | 6 | 39 |
| 78 | 194443 | S1200 | Q12 | M | 26-30 | 26 | 32 | 39 | 23 | 42 |
| 79 | 196851 | S1200 | Q09 | M | 26-30 | 31 | 23 | 30 | 17 | 33 |
| 80 | 196952 | S1200 | Q13 | F | 26-30 | 34 | 13 | 42 | 14 | 24 |
| 81 | 198047 | S1200 | Q13 | F | 26-30 | 31 | 38 | 37 | 20 | 35 |
| 82 | 199352 | S1200 | Q12 | F | 22-25 | 27 | 28 | 41 | 14 | 32 |
| 83 | 200513 | S1200 | Q12 | M | 22-25 | 33 | 32 | 34 | 19 | 33 |
| 84 | 204218 | S1200 | Q09 | M | 26-30 | 27 | 31 | 37 | 12 | 40 |
| 85 | 206323 | S1200 | Q12 | F | 31-35 | 34 | 26 | 35 | 9 | 31 |
| 86 | 206525 | S1200 | Q09 | M | 26-30 | 19 | 31 | 40 | 3 | 37 |
| 87 | 206727 | S1200 | Q09 | M | 22-25 | 25 | 21 | 28 | 27 | 30 |
| 88 | 206828 | S1200 | Q11 | M | 22-25 | 31 | 28 | 47 | 4 | 32 |
| 89 | 206929 | S1200 | Q12 | M | 31-35 | 33 | 26 | 35 | 11 | 34 |
| 90 | 208630 | S1200 | Q12 | M | 31-35 | 26 | 24 | 36 | 16 | 26 |
| 91 | 210112 | S1200 | Q11 | M | 22-25 | 26 | 41 | 27 | 16 | 36 |
| 92 | 211619 | S1200 | Q13 | M | 26-30 | 31 | 32 | 37 | 9 | 24 |
| 93 | 211821 | S1200 | Q06 | M | 26-30 | 32 | 28 | 32 | 20 | 24 |
| 94 | 213017 | S1200 | Q11 | M | 22-25 | 35 | 30 | 35 | 12 | 34 |
| 95 | 213522 | S1200 | Q12 | M | 26-30 | 27 | 25 | 27 | 27 | 10 |
| 96 | 219231 | S1200 | Q05 | M | 26-30 | 37 | 37 | 26 | 29 | 24 |
| 97 | 227533 | S1200 | Q08 | F | 31-35 | 41 | 29 | 43 | 6 | 40 |
| 98 | 238033 | S1200 | Q09 | M | 31-35 | 30 | 40 | 23 | 24 | 23 |
| 99 | 239136 | S1200 | Q09 | F | 26-30 | 40 | 36 | 44 | 15 | 34 |
| 100 | 248238 | S1200 | Q13 | F | 31-35 | 37 | 31 | 36 | 18 | 12 |
| 101 | 255740 | S1200 | Q10 | F | 26-30 | 35 | 24 | 37 | 34 | 29 |
| 102 | 257946 | S1200 | Q08 | F | 26-30 | 29 | 28 | 38 | 20 | 31 |
| 103 | 274542 | S1200 | Q13 | F | 26-30 | 31 | 32 | 38 | 25 | 37 |
| 104 | 281135 | S1200 | Q12 | M | 26-30 | 35 | 21 | 39 | 11 | 30 |
| 105 | 286347 | S1200 | Q13 | F | 26-30 | 45 | 35 | 25 | 27 | 28 |
| 106 | 299760 | S1200 | Q09 | M | 26-30 | 34 | 38 | 25 | 19 | 30 |
| 107 | 300719 | S1200 | Q13 | M | 26-30 | 32 | 31 | 34 | 14 | 26 |
| 108 | 314225 | S1200 | Q12 | M | 22-25 | 28 | 28 | 30 | 19 | 32 |
| 109 | 325129 | S1200 | Q13 | M | 31-35 | 38 | 31 | 33 | 24 | 29 |
| 110 | 329844 | S1200 | Q07 | M | 26-30 | 33 | 26 | 39 | 13 | 36 |
| 111 | 342129 | S1200 | Q09 | M | 26-30 | 39 | 30 | 37 | 9 | 42 |
| 112 | 349244 | S1200 | Q08 | M | 26-30 | 29 | 26 | 40 | 14 | 23 |
| 113 | 350330 | S1200 | Q13 | F | 22-25 | 37 | 32 | 32 | 8 | 32 |
| 114 | 360030 | S1200 | Q13 | F | 31-35 | 37 | 22 | 35 | 20 | 26 |
| 115 | 368551 | S1200 | Q09 | M | 26-30 | 26 | 34 | 39 | 7 | 35 |
| 116 | 368753 | S1200 | Q12 | F | 31-35 | 36 | 21 | 43 | 10 | 19 |
| 117 | 376247 | S1200 | Q13 | F | 26-30 | 36 | 24 | 38 | 24 | 32 |
| 118 | 378756 | S1200 | Q12 | M | 26-30 | 36 | 36 | 29 | 25 | 30 |
| 119 | 385046 | S1200 | Q09 | M | 26-30 | 32 | 30 | 37 | 14 | 28 |
| 120 | 392750 | S1200 | Q08 | M | 31-35 | 34 | 27 | 27 | 20 | 22 |
| 121 | 394956 | S1200 | Q12 | F | 26-30 | 38 | 29 | 38 | 15 | 37 |
| 122 | 401422 | S1200 | Q08 | F | 31-35 | 42 | 29 | 33 | 12 | 38 |
| 123 | 413934 | S1200 | Q09 | F | 31-35 | 28 | 35 | 34 | 28 | 22 |
| 124 | 419239 | S1200 | Q12 | F | 26-30 | 33 | 26 | 41 | 21 | 26 |
| 125 | 421226 | S1200 | Q13 | M | 31-35 | 34 | 29 | 27 | 18 | 31 |
| 126 | 454140 | S1200 | Q12 | M | 31-35 | 30 | 28 | 35 | 20 | 31 |
| 127 | 461743 | S1200 | Q11 | M | 26-30 | 25 | 25 | 37 | 10 | 33 |
| 128 | 462139 | S1200 | Q13 | F | 26-30 | 41 | 32 | 44 | 5 | 37 |
| 129 | 463040 | S1200 | Q13 | F | 31-35 | 37 | 28 | 26 | 17 | 27 |
| 130 | 469961 | S1200 | Q13 | F | 26-30 | 34 | 28 | 35 | 14 | 26 |
| 131 | 481042 | S1200 | Q12 | F | 31-35 | 32 | 24 | 36 | 18 | 27 |
| 132 | 510225 | S1200 | Q08 | F | 26-30 | 39 | 45 | 32 | 26 | 25 |
| 133 | 513130 | S1200 | Q12 | F | 26-30 | 41 | 22 | 33 | 9 | 38 |
| 134 | 516742 | S1200 | Q11 | M | 31-35 | 30 | 26 | 45 | 3 | 36 |
| 135 | 518746 | S1200 | Q12 | M | 22-25 | 31 | 18 | 33 | 19 | 30 |
| 136 | 519647 | S1200 | Q13 | M | 26-30 | 22 | 20 | 40 | 15 | 20 |
| 137 | 531940 | S1200 | Q13 | F | 22-25 | 39 | 44 | 31 | 29 | 36 |
| 138 | 541640 | S1200 | Q06 | M | 26-30 | 39 | 47 | 30 | 12 | 36 |
| 139 | 550439 | S1200 | Q12 | M | 22-25 | 30 | 24 | 35 | 22 | 35 |
| 140 | 552241 | S1200 | Q12 | M | 22-25 | 36 | 24 | 36 | 15 | 29 |
| 141 | 555954 | S1200 | Q09 | M | 26-30 | 31 | 28 | 34 | 14 | 30 |
| 142 | 558657 | S1200 | Q13 | M | 22-25 | 34 | 32 | 33 | 28 | 24 |
| 143 | 558960 | S1200 | Q12 | F | 31-35 | 41 | 30 | 31 | 25 | 33 |
| 144 | 559457 | S1200 | Q13 | M | 31-35 | 34 | 28 | 36 | 14 | 23 |
| 145 | 561949 | S1200 | Q12 | M | 26-30 | 27 | 14 | 32 | 16 | 28 |
| 146 | 567759 | S1200 | Q09 | F | 26-30 | 32 | 29 | 28 | 13 | 28 |
| 147 | 569965 | S1200 | Q13 | F | 31-35 | 32 | 38 | 27 | 20 | 33 |
| 148 | 578057 | S1200 | Q13 | F | 26-30 | 42 | 24 | 42 | 16 | 34 |
| 149 | 578158 | S1200 | Q13 | M | 26-30 | 33 | 42 | 43 | 7 | 32 |
| 150 | 589567 | S1200 | Q13 | M | 22-25 | 25 | 29 | 39 | 18 | 35 |
| 151 | 590047 | S1200 | Q09 | M | 22-25 | 29 | 36 | 21 | 25 | 29 |
| 152 | 615441 | S1200 | Q12 | M | 22-25 | 34 | 22 | 35 | 9 | 27 |
| 153 | 623137 | S1200 | Q13 | F | 31-35 | 25 | 31 | 19 | 15 | 31 |
| 154 | 634748 | S1200 | Q13 | F | 22-25 | 43 | 40 | 40 | 20 | 33 |
| 155 | 635245 | S1200 | Q12 | M | 26-30 | 27 | 32 | 29 | 4 | 42 |
| 156 | 644246 | S1200 | Q12 | F | 31-35 | 37 | 25 | 39 | 15 | 32 |
| 157 | 654552 | S1200 | Q12 | F | 31-35 | 34 | 26 | 35 | 18 | 33 |
| 158 | 675661 | S1200 | Q12 | M | 31-35 | 24 | 30 | 30 | 31 | 25 |
| 159 | 680452 | S1200 | Q13 | F | 31-35 | 39 | 17 | 32 | 19 | 38 |
| 160 | 688569 | S1200 | Q02 | F | 26-30 | 34 | 18 | 42 | 32 | 21 |
| 161 | 692964 | S1200 | Q12 | M | 22-25 | 25 | 33 | 35 | 28 | 38 |
| 162 | 694362 | S1200 | Q13 | F | 22-25 | 37 | 26 | 37 | 22 | 32 |
| 163 | 698168 | S1200 | Q12 | M | 22-25 | 43 | 39 | 28 | 23 | 38 |
| 164 | 701535 | S1200 | Q13 | F | 31-35 | 36 | 28 | 35 | 7 | 35 |
| 165 | 723141 | S1200 | Q12 | M | 31-35 | 33 | 39 | 36 | 31 | 29 |
| 166 | 728454 | S1200 | Q12 | M | 22-25 | 27 | 22 | 39 | 18 | 32 |
| 167 | 760551 | S1200 | Q12 | F | 26-30 | 38 | 25 | 40 | 18 | 30 |
| 168 | 763557 | S1200 | Q09 | M | 26-30 | 28 | 37 | 27 | 27 | 33 |
| 169 | 765864 | S1200 | Q08 | F | 26-30 | 35 | 35 | 26 | 27 | 33 |
| 170 | 774663 | S1200 | Q13 | M | 26-30 | 27 | 31 | 40 | 10 | 42 |
| 171 | 788674 | S1200 | Q11 | F | 31-35 | 41 | 18 | 35 | 23 | 35 |
| 172 | 809252 | S1200 | Q11 | F | 26-30 | 28 | 24 | 33 | 15 | 35 |
| 173 | 814548 | S1200 | Q13 | F | 31-35 | 34 | 31 | 43 | 13 | 33 |
| 174 | 815247 | S1200 | Q08 | M | 26-30 | 30 | 15 | 39 | 11 | 30 |
| 175 | 818455 | S1200 | Q08 | M | 31-35 | 38 | 40 | 24 | 17 | 31 |
| 176 | 822244 | S1200 | Q08 | F | 26-30 | 39 | 23 | 41 | 16 | 39 |
| 177 | 825553 | S1200 | Q09 | M | 26-30 | 33 | 23 | 29 | 20 | 33 |
| 178 | 825654 | S1200 | Q12 | F | 31-35 | 42 | 21 | 48 | 5 | 37 |
| 179 | 827052 | S1200 | Q13 | F | 26-30 | 35 | 32 | 35 | 16 | 24 |
| 180 | 828862 | S1200 | Q13 | F | 26-30 | 44 | 19 | 40 | 14 | 25 |
| 181 | 832651 | S1200 | Q12 | M | 22-25 | 33 | 31 | 23 | 10 | 34 |
| 182 | 869472 | S1200 | Q11 | F | 22-25 | 37 | 30 | 34 | 14 | 34 |
| 183 | 878877 | S1200 | Q09 | M | 26-30 | 30 | 28 | 32 | 13 | 30 |
| 184 | 884064 | S1200 | Q07 | F | 31-35 | 34 | 26 | 38 | 29 | 22 |
| 185 | 886674 | S1200 | Q11 | M | 22-25 | 26 | 37 | 23 | 22 | 23 |
| 186 | 888678 | S1200 | Q13 | M | 26-30 | 27 | 33 | 41 | 5 | 28 |
| 187 | 908860 | S1200 | Q10 | M | 26-30 | 32 | 40 | 11 | 34 | 24 |
| 188 | 911849 | S1200 | Q13 | F | 31-35 | 39 | 43 | 31 | 14 | 37 |
| 189 | 929464 | S1200 | Q11 | F | 26-30 | 34 | 39 | 33 | 34 | 32 |
| 190 | 933253 | S1200 | Q12 | F | 26-30 | 31 | 28 | 39 | 12 | 28 |
| 191 | 943862 | S1200 | Q09 | M | 26-30 | 37 | 29 | 33 | 10 | 29 |
| 192 | 969476 | S1200 | Q12 | M | 31-35 | 26 | 21 | 35 | 20 | 19 |
| 193 | 970764 | S1200 | Q09 | F | 22-25 | 41 | 30 | 36 | 17 | 36 |
| 194 | 971160 | S1200 | Q10 | M | 26-30 | 32 | 36 | 37 | 30 | 34 |
| 195 | 973770 | S1200 | Q13 | M | 22-25 | 28 | 27 | 36 | 27 | 26 |
| 196 | 987074 | S1200 | Q10 | M | 22-25 | 37 | 28 | 28 | 20 | 24 |
| 197 | 989987 | S1200 | Q13 | M | 31-35 | 35 | 24 | 34 | 17 | 30 |
| 198 | 995174 | S1200 | Q13 | M | 22-25 | 27 | 30 | 41 | 18 | 36 |

**Table S2.** The WU-Minn HCP Retest Data is used to evaluate the robustness of our method to construct individual myelin network. Here, twenty participants are included in our study. The T1-weighted and T2-weighted data were processed using the MRTool (https://www.nitrc.org/projects/mrtool, version 1.4.2) implemented in the SPM12.

|  | SUBJECT | RELEASE | ACQUISITION | GENDER | AGE |
| --- | --- | --- | --- | --- | --- |
| 1 | 105923 | MEG2 | Q07 | F | 31-35 |
| 2 | 122317 | Q3 | Q04 | M | 31-35 |
| 3 | 137128 | Q1 | Q02 | F | 31-35 |
| 4 | 139839 | S900 | Q07 | M | 26-30 |
| 5 | 149337 | Q1 | Q02 | M | 31-35 |
| 6 | 149741 | S500 | Q05 | M | 26-30 |
| 7 | 158035 | Q2 | Q02 | F | 26-30 |
| 8 | 169343 | Q2 | Q02 | F | 31-35 |
| 9 | 172332 | Q1 | Q02 | F | 26-30 |
| 10 | 177746 | Q2 | Q02 | F | 26-30 |
| 11 | 192439 | Q1 | Q01 | F | 31-35 |
| 12 | 194140 | Q1 | Q01 | F | 26-30 |
| 13 | 195041 | S500 | Q07 | F | 31-35 |
| 14 | 200109 | S500 | Q06 | F | 31-35 |
| 15 | 204521 | S500 | Q07 | F | 31-35 |
| 16 | 287248 | MEG2 | Q08 | F | 26-30 |
| 17 | 433839 | S500 | Q04 | M | 26-30 |
| 18 | 627549 | Q2 | Q02 | F | 31-35 |
| 19 | 660951 | MEG2 | Q08 | F | 26-30 |
| 20 | 877168 | Q1 | Q01 | F | 31-35 |

## Notes

### Competing Interest Statement

The authors have declared no competing interest.

